# Bright, fluorogenic and photostable avidity probes for RNA imaging

**DOI:** 10.1101/2021.11.02.466936

**Authors:** Bastian Bühler, Anja Benderoth, Daniel Englert, Franziska Grün, Janin Schokolowski, Andres Jäschke, Murat Sunbul

## Abstract

Fluorescent light-up aptamers (FLAPs) emerged as valuable tools to visualize RNA, but are mostly limited by poor brightness, low photostability and high fluorescence background. In this study, we combine bivalent (silicon) rhodamine fluorophores with dimeric FLAPs to yield bright and photostable complexes with low picomolar dissociation constants. Our avidity-based approach resulted in extreme binding strength and high fluorogenicity, enabling single mRNA tracking in living cells.

## Main

Genetically encoded fluorescent light-up aptamers (FLAPs) as imaging modalities have recently started to provide insights into myriad functions of RNA^1-4^. However, the use of most FLAPs for single-molecule RNA tracking is still limited due to poor brightness, rapid photobleaching and weak binding affinity of the aptamer:dye complexes, in addition to low cellular permeability and fluorogenicity of the dyes^4-6^. To overcome these limitations, we report here a system consisting of synthetic aptamer dimers and bivalent state-of-the-art fluorophores. As FLAPs, we choose RhoBAST and SiRA, two bright and photostable (silicon) rhodamine-binding aptamers, that enabled SMLM and STED imaging of RNA in living cells for the first time^1, 7^. The dimeric system we introduced in this study mimics the interactions between bivalent antibodies and multivalent antigens (Figure 1a, b). Exploiting the avidity concept, we created fluorogenic aptamer-dye pairs with the strongest binding to date, enabling tracking of RNA molecules in living cells with a high signal-to-background ratio.

**Figure 1.**
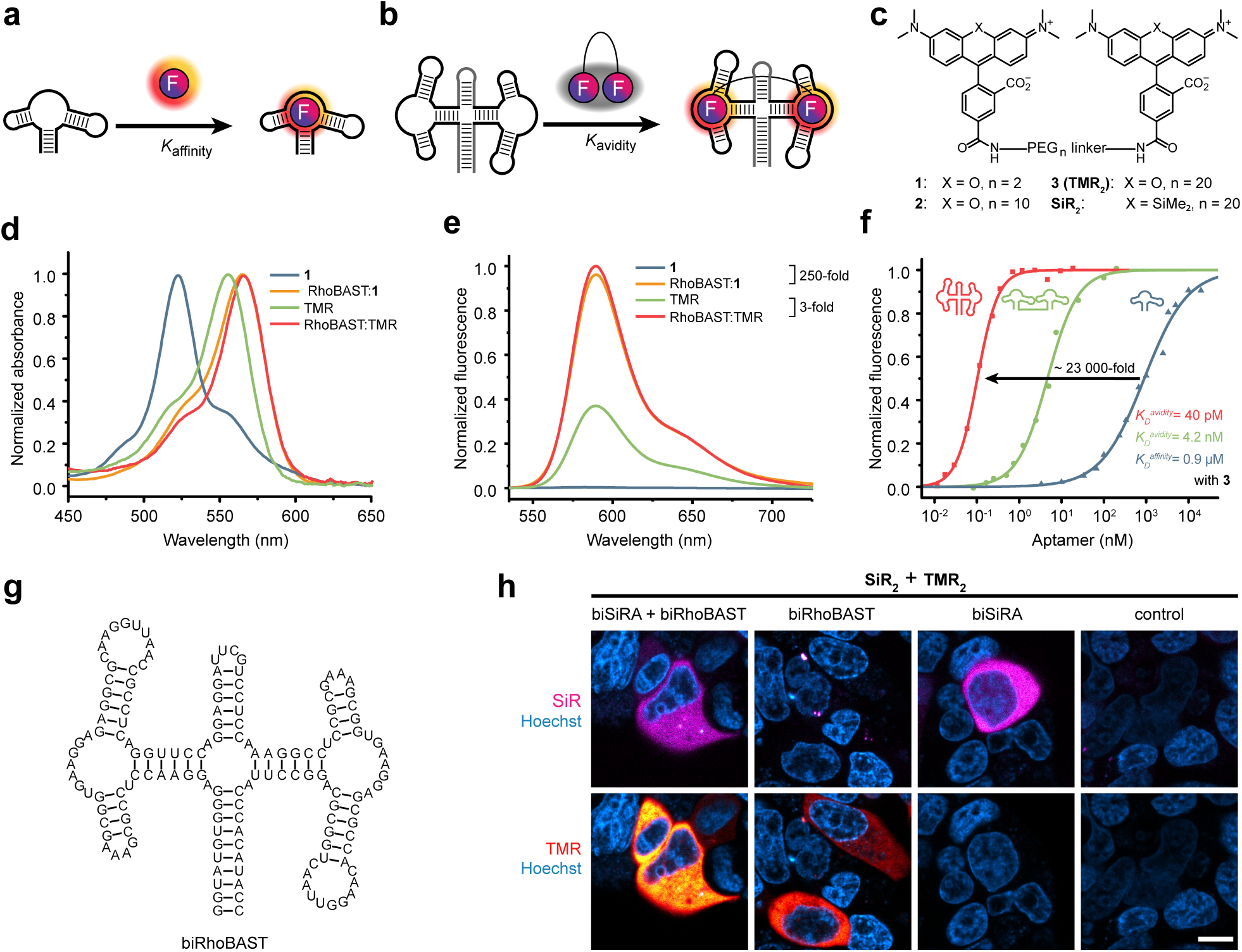
Avidity-based fluorogenic RNA imaging. **a)** Binding of an aptamer to a fluorophore with a certain affinity. **b)** Avidity-based binding of a dimeric aptamer to a fluorogenic bivalent fluorophore. Unbound bivalent fluorophores are non-fluorescent due to H-dimer formation. **c)** Bivalent fluorophores synthesized in this study. Two (silicon) rhodamine fluorophores were conjugated via polyethylene glycol (PEG) linkers with different lengths. **d)** Absorption and **e)** emission spectra of bivalent TMR (**1**) in comparison to monomeric TMR. **1** showed a characteristic blue-shifted H-dimer absorption maximum which largely disappeared upon binding to RhoBAST. **f)** Fluorescence increase upon binding of **3** (**TMR**_**2**_) to monomeric and dimeric (flexible and rigid) RhoBAST. Due to avidity effect, dissociation constant of biRhoBAST: **TMR**_**2**_ (*K*_D_^avidity^ = 40 pM) is ∼23 000-fold lower than that of RhoBAST:**TMR**_**2**_. **g)** Predicted secondary structure of biRhoBAST consisting of a four-way junction with a stabilizing stem loop and two synonymous aptamer units. **h)** Orthogonal labeling of circular biSiRA and biRhoBAST aptamers with **SiR**_**2**_ (50 nM) and **TMR**_**2**_ (50 nM) in live HEK293T cells. Scale bar, 10 µm.

First, we synthesized a series of bivalent TMR dyes (Figure 1c) with polyethylene glycol linkers (*n* = 2, 10, 20 ethylene glycol units; designated as **1, 2**, and **3**, respectively). All bivalent TMR dyes were essentially non-fluorescent and displayed blue-shifted absorption band in aqueous solution compared to monomeric TMR (Figure 1d, Supplementary Figure 1), suggesting intramolecular H-dimer formation^8^. We validated the intramolecular nature of the dimerization by measuring the fluorescence of the bivalent TMR dyes as a function of concentration in the low nanomolar (nM) range (Supplementary Figure 1f). The observed excitation and emission maxima of the bivalent TMR dyes were similar to those of monomeric TMR (Supplementary Figure 1g), indicating that the low residual fluorescence is due to the dynamic equilibrium between the fluorescent form and the non-fluorescent H-dimer form (Supplementary Figure 1a)^9, 10^. As expected, the proportion of the H-dimer form increases with decreasing linker length due to increased local dye concentration, consistent with the observed fluorescence intensities of **1** < **2** < **3** (Supplementary Figure 1f). The strength of the H-dimer formation prevents significant alteration of the absorption spectra or the fluorescence intensity in the presence of detergent, proteins or total RNA (Supplementary Figure 1h,i). Therefore, bivalent TMR dyes should exhibit low unspecific fluorescence background in living cells.

Next, we demonstrated that the RhoBAST aptamer can bind bivalent TMR dyes and disrupt H-dimers, resulting in a red-shift of the absorption maximum and an increase in fluorescence intensities of **1, 2** and **3** up to 250-fold (Figure 1d,e). In comparison, RhoBAST yielded only a ∼3-fold fluorescence turn-on with monomeric TMR (Figure 1e). We also determined dissociation constants (*K*_D_ ^mono^) of RhoBAST complexes with bivalent dyes **1, 2** and **3** as >46, 1.8 and 0.9 µM, respectively, again showing a strong dependence on linker length (Supplementary Table 1). The observed *K*_D_^mono^ values are several orders of magnitude higher compared to RhoBAST:TMR (*K*_D_ = 15 nM)^1^, emphasizing the high stability of TMR H-dimers (Figure 1f, Supplementary Figure 2).

**Figure 2.**
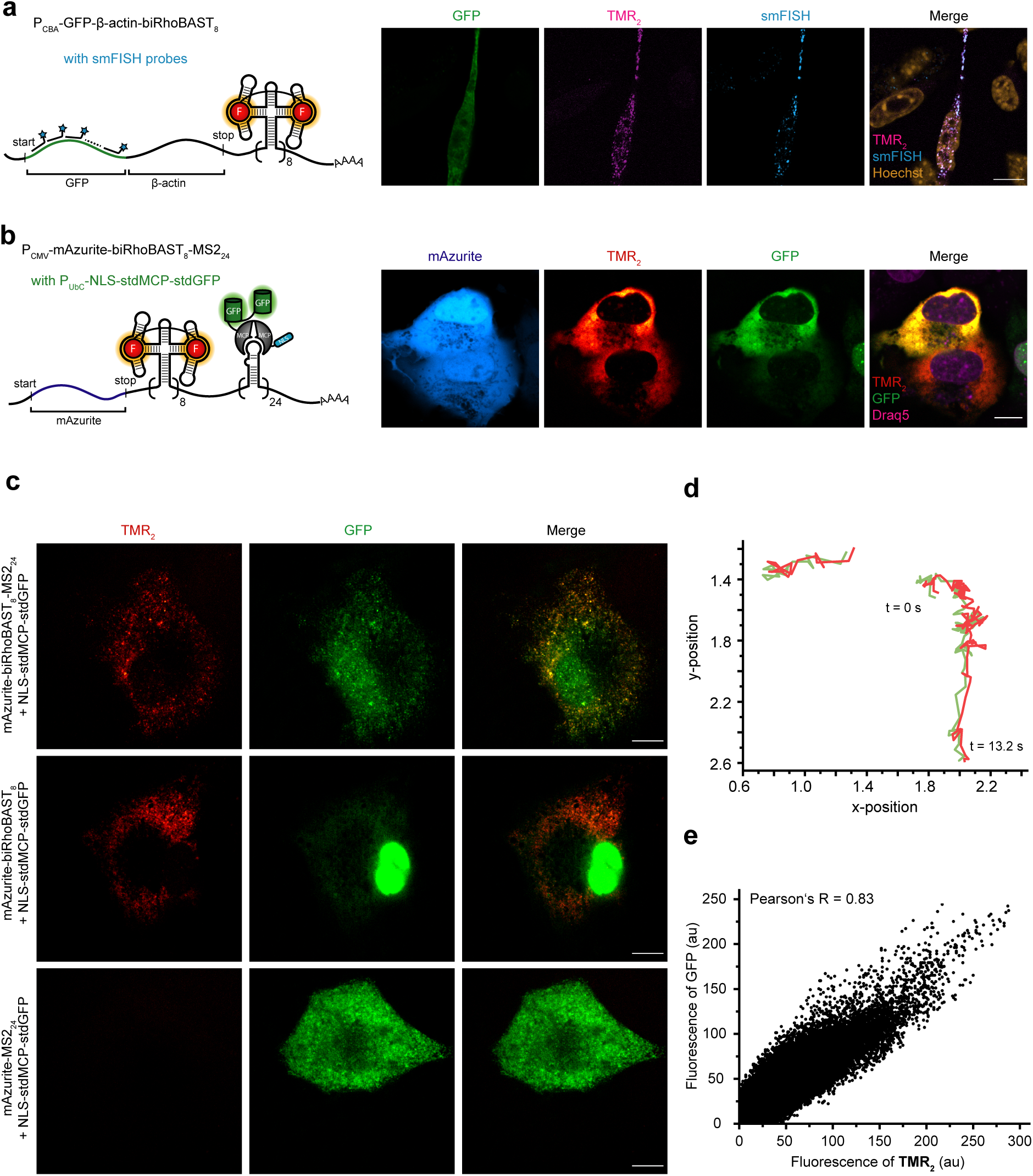
The biRhoBAST:TMR_2_ system enables single molecule RNA tracking in live cells. **a)** Confocal images of fixed MEF cells expressing *GFP*-β-*actin-biRhoBAST*_*8*_ mRNA after incubation with smFISH probes (targeting the GFP sequence) in the presence of **TMR**_**2**_ (50 nM). The overlay image shows significant colocalization of smFISH and TMR signals. Scale bar, 5 µm. **b)** Confocal images of live Cos7 cells co-transfected with P_CMV_- mAzurite-biRhoBAST_8_-MS2_24_ and P_UbC_-NLS-stdMCP-stdGFP plasmids in the presence of 50 nM **TMR**_**2**_. The overlay image shows significant co-localization of GFP and TMR signals. Scale bar, 10 µm. **c)** Spinning disc confocal images of Cos7 cells expressing low levels of MCP-GFP and *mAzurite-biRhoBAST*_*8*_*-MS2*_*24*_ (top), *mAzurite-biRhoBAST*_*8*_ (middle), *mAzurite-MS2*_*24*_ (bottom) in the presence of 500 nM **TMR**_**2**_. Single mRNA molecules were visualized with a frame rate of 10 s^−1^. Scale bars, 10 µm. **d)** Extracted trajectories at different time points of single RNA tracks in the image shown in panel **c** (top), revealing synchronous movements of **TMR**_**2**_ and GFP signals over time. **e)** Colocalization test of the cytosolic **TMR**_**2**_ and GFP signals in the image shown in panel **c** (top), resulting in a Pearson’s R value of 0.83.

To improve the binding affinity between the aptamer and the bivalent TMR dyes, we aimed to apply the avidity concept^11, 12^. We designed a dimeric version of RhoBAST by linking two synonymous aptamers with a flexible 10-nt linker sequence and determined the dissociation constants of its complexes with the bivalent TMR dyes (Supplementary Figure 2). For **1, 2** and **3** we measured values of 49, 6.0 and 4.2 nM, respectively, revealing >940-, 300- and 220-fold improved binding strength compared to their respective *K*_D_^mono^. Next, to reduce the entropic penalty and enhance the avidity effect, we introduced a scaffold that locks the orientation and distance between the two RhoBAST units (biRhoBAST, Figure 1g). Indeed, **1, 2** and **3** bind biRhoBAST >4.9×10^5^-, 3.0×10^4^- and 2.3×10^4^-fold stronger compared to monomeric RhoBAST, displaying *K*_D_ ^avidity^ values of 93, 61 and 40 pM, which are the lowest reported *K*_D_ values for RNA:fluorophore complexes (Figure 1f, Supplementary Figure 2, Supplementary Table 1). Furthermore, as shown by the absorption spectra, binding of biRhoBAST effectively disrupted TMR H-dimers in **1, 2** and **3** and resulted in 284-, 92- and 55-fold fluorescence turn-on, respectively (Supplementary Figure 3).

Next, we investigated the performance of **1, 2** and **3** in live cells. Confocal images of HEK293T cells expressing circular biRhoBAST aptamer^13^ were acquired in the presence of these fluorogenic dyes. In live cells, **3** evidently outcompeted **1** and **2** with higher signal-to background ratios (Supplementary Figure 4), which prompted us to characterize the complex between **3** (designated as **TMR**_**2**_ from now on) and biRhoBAST in more detail. The biRhoBAST:**TMR**_**2**_ complex has an excellent quantum yield (QY = 0.88) and a high extinction coefficient (165 000 M^-1^cm^-1^), resulting in an exceptionally bright FLAP (Supplementary Table 2,4).

The fluorescence of the complex was neither sensitive towards magnesium ion concentration within the biologically relevant window, nor dependent on monovalent cations, and showed a high thermal stability (Supplementary Figure 5a-c). Due to the avidity effect, biRhoBAST:**TMR**_**2**_ displays an extremely slow dissociation (*k*_off_ = 3.8×10^−4^ s^−1^), ∼10^4^-fold slower than monomeric RhoBAST:TMR-DN (*k*_off_ = 2.9 s^−1^)^1^(Supplementary Figure 6). biRhoBAST:**TMR**_**2**_ is also remarkably photostable. Compared to Pepper:HBC620, reported to be more photo-stable than most FLAPs^5^, we did not observe significant photo-bleaching of biRhoBAST:**TMR**_**2**_ *in vitro* (Supplementary Figure 7). Besides Pepper:HBC620, we also compared the photostability of our novel system to Broccoli:BI and Mango2:TO1-biotin, both of which have recently been used to image single RNA molecules^14, 15^. Under constant illumination in live cells, the fluorescence intensities of Mango2:TO1-biotin, Broccoli:BI and Pepper:HBC620 decreased to 50 % of their initial value after 0.3 s, 1.1 s and 10.3 s, respectively. In contrast, the fluorescence of biRhoBAST:**TMR**_**2**_ was reduced by only 3 % after 90 s illumination, indicating excellent photostability (Supplementary Figure 7).

Next, we asked if we could apply the same avidity-based concept to improve the properties of the SiRA system, in which fluorogenicity arises from spirolactonization of the fluorophore (Supplementary Figure 8a). We synthesized a bivalent silicon rhodamine dye (**SiR**_**2**_) and designed the dimeric aptamer biSiRA (Supplementary Figure 8b-d). The biSiRA:**SiR**_**2**_ system showed a much higher fluorogenicity (100-fold turn-on) and extremely improved affinity (*K*_D_^avidity^ = 290 pM) compared to SiRA:SiR (7-fold turn-on and *K*_D_ of 430 nM) (Supplementary Figure 8e-i)^7^. In the biSiRA:**SiR**_**2**_ system, the origin of the fluorogenicity was found to be a result of H-dimerization as well as spirolactonization (Supplementary Note 1). biSiRA:**SiR**_**2**_ is thermostable and its fluorescence does not depend on monovalent cations, but is strongly affected by magnesium concentration (Supplementary Figure 9a-f). Because of its excellent fluorescence quantum yield (QY = 0.98) and high extinction coefficient (200 000 M^- 1^cm^-1^), biSiRA:**SiR**_**2**_ is the brightest FLAP reported to date.

The **SiR**_**2**_ avidity probe enabled confocal imaging of circular biSiRA in live cells, which was not possible with sufficient signal-to-background ratios using the original SiRA system (Supplementary Figure 10). Moreover, we verified the orthogonality of biRhoBAST:**TMR**_**2**_ and biSiRA:**SiR**_**2**_ systems by simultaneously imaging two RNAs (Figure 1h). Similar to biRhoBAST:**TMR**_**2**_, biSiRA:**SiR**_**2**_ also demonstrated excellent resistance against photobleaching compared to other reported FLAPs (Supplementary Figure 7).

Next, we cloned the biAptamers into the 3′ untranslated region (UTR) of *mEGFP* and β-*actin* mRNAs transcribed from different promoters. **TMR**_**2**_ enabled imaging of target mRNAs with only a single biRhoBAST tag where the contact-quenched TMR-DN probe failed due to a much lower signal-to-background ratio (Supplementary Figure 11). On the other hand, **SiR**_**2**_ did not yield adequate signal-to-background ratios for imaging these mRNAs with a single biSiRA tag, probably due to the complex environment in live cells and biSiRA’s strong dependence on magnesium ions (Supplementary Figure 9f and Supplementary Figure 10c,d). We therefore focused on biRhoBAST:**TMR**_**2**_ for single-molecule imaging experiments and cloned multiple synonymous biRhoBAST repeats into the 3′ UTR of *mEGFP* mRNA to further increase the signal-to-background ratio. We verified that TMR fluorescence in living cells increases almost linearly with the numbers of repeating biRhoBAST units (Supplementary Figure 12). Then, we imaged *GFP*-β-*actin* mRNA carrying a zip code and eight repeats of biRhoBAST in the 3′ UTR in fixed fibroblasts using **TMR**_**2**_ and single-molecule fluorescence in-situ hybridization (smFISH) probes targeting the GFP sequence^16, 17^. Substantial colocalization of smFISH and **TMR**_**2**_ probes with a Pearson’s R of 0.91 validates that; (i) the interaction between biRhoBAST and **TMR**_**2**_ is highly specific and sensitive; (ii) biRhoBAST can still bind to **TMR**_**2**_ after formaldehyde fixation; (iii) the biRhoBAST:**TMR**_**2**_ system enables imaging of full-length RNA transcripts and not just decay fragments (Figure 2a and Supplementary Figure 13).

After showing specific and full-length *GFP*-β-*actin* mRNA imaging in fixed cells, we created a new construct to transcribe *mAzurite* mRNA fused to 8 and 24 repeats of biRhoBAST and MS2, respectively, at the 3′ UTR. This way we can compare biRhoBAST:**TMR**_**2**_ to the MS2:MCP-GFP (MCP; MS2 coat protein) system, a gold standard for single molecule RNA tracking in live cells^18, 19^. Cells co-expressing *mAzurite*-*biRhoBAST*_8_*-MS2*_24_ mRNA and MCP-GFP protein were incubated with **TMR**_**2**_ and imaged. As expected, images showed significant colocalization of GFP and TMR signals (Figure 2b). Moreover, biRhoBAST:**TMR**_**2**_ showed a higher signal-to-background ratio compared to the MS2 system, despite containing fewer fluorophores (16 TMR versus 48 GFP) (Supplementary Figure 14a,b). biRhoBAST:**TMR**_**2**_ enabled the visualization of RNAs localized in the nucleus, which was not possible with the standard MS2 system (Supplementary Figure 14c). When *mAzurite*-*biRhoBAST*_8_-*MS2*_*2*4_ mRNA expression was low, we could detect highly mobile puncta in the cytosol using a spinning disc confocal microscope (Figure 2c and Supplementary Video 1). Time-resolved extracted trajectories of the single particles showed synchronous movements of GFP and TMR signals (Figure 2d) and analysis of all TMR and GFP signals in the cytosol indicated significant colocalization (Pearson’s R of 0.83) (Figure 2e). These data demonstrate that biRhoBAST:**TMR**_**2**_ enables visualization of single RNA molecules in live cells with a high signal-to-background ratio. Furthermore, by plotting the mean square displacement (MSD) versus the delay time of several distinct *mAzurite*-*biRhoBAST*_*8*_ mRNAs in live cells, we calculated the diffusion coefficient (D) of mobile mRNAs to be 0.116 µm^2^s^-1^, which is in consistent with those of *mAzurite*-*MS2*_*2*4_ mRNAs (Supplementary Figure 15 and Supplementary Video 2 and 3) and values reported in the literature^20^.

In summary, we have demonstrated a general avidity-based approach to engineer highly fluorogenic and extreme-affinity aptamer:fluorophore pairs by using bivalent state-of-the-art fluorophores and dimeric fluorophore-binding aptamers. The lowest reported *K*_D_ between a fluorophore and aptamer as well as the outstanding brightness enabled orthogonal imaging of red and near-infrared labelled RNA in live cells. Additionally, we were able to track single mRNAs with our novel tag. In the future, we aim to investigate the lifecycle of RNA at the molecular level using these orthogonal dual-color probes.

## Author Contributions and Notes

B.B, A.J. and M.S. designed the study. B.B. and A.B. performed synthesis. B.B., A.B., D.E., F.G. and J.S. performed *in vitro* characterizations. B.B. created all plasmid constructs and conducted live cell confocal imaging and single RNA tracking. B.B wrote the initial draft of the manuscript and all authors participated in revising and editing.

The authors declare no conflict of interest. This article contains supporting information online.

## Acknowledgments

M.S. and A.J. were supported by the Deutsche Forschungsgemeinschaft (DFG grant no. Ja794/11). We thank the Nikon Imaging Center, Heidelberg for granting access to their facilities and U. Engel for technical advice in fluorescence microscopy.

## Materials and Methods

### General

All used reagents were purchased from Thermo Fisher Scientific, Iris Biotech, Sigma-Aldrich, Alfa Aesar GmbH or TCI Chemicals and were used without further purification, unless otherwise stated. Deuterated solvents for NMR spectroscopy (DMSO*-d*6, CDCl3 and MeOH-*d*4)), were purchased from Euriso-Top GmbH. All NMR spectra were recorded on a Varian Mercury Plus 300 MHz spectrometer or a Varian 500 MHz NMR spectrometer. Peak shifts were reported relative to the solvent peaks. Chemical shifts are given in parts per million (ppm) and the coupling constants in Hertz (Hz). Multiplicities are given as shift of the centre signal using the following abbreviations: s (singlet), d (doublet), t (triplet), q (quartet), m (multiplet), dd (doublet of doublet), dt (doublet of triplet). MestReNova 9.0.1 developed by MESTRELAB RESEARCH was used for NMR data processing. Polygram Sil Silica gel from Fluka with a pore size of 60° and a particle size range of 40-63 μm was used for normal phase column chromatography. The solvents were *p*.*a*. quality. The fluorophores for photophysical measurements or biological applications were purified via reverse-phase semi-preparative HPLC and afterwards their purity was verified via analytical HPLC. We used a 1100 HPLC from Agilent Technologies, *Inc* and Luna 5 µm C18(2) 100° column, 250 mm × 10 mm (Phenomenex). Compounds were eluted using gradients of water (H_2_O containing 0.1 % trifluoroacetic acid) and acetonitrile (ACN containing 0.1 % trifluoroacetic acid) with a constant flow rate of 6 mL/min. For detailed purification gradients see experimental part (Supplementary Note 2). All HPLC fractions were dried by lyophilization (Christ). High resolution mass spectrometry (HRMS) was performed on a Bruker microTOF-QII mass spectrometer. Fluorescence measurements were performed on a Jasco Spectrofluorometer FP-6500 or FP-8500. Absorption spectra were recorded on a Cary 50 UV. Agarose gels were stained with ethidium bromide and visualized by UV illumination using an AlphaImager TM 2200 (Alpha Innotech). DNA and RNA concentrations were determined using a NanoDrop One (Thermo Fisher Scientific) spectrophotometer. All oligonucleotides were purchased from Integrated DNA Technologies (IDT). Plasmids were obtained from Addgene.

Bacterial strains (DH5α) were purchased from Thermo Fisher Scientific and grown at 37°C with shaking at 160 rpm in Luria-Bertani (LB) medium. Restriction enzyme cloning was performed with Thermo Fisher Scientific FastDigest® enzymes according to the manufacturer’s instructions. Sanger sequencing was carried out at Microsynth Seqlab with standard primers, if not stated otherwise.

6xASBT buffer contains 120 mM HEPES pH 7.4, 750 mM KCl, 30 mM MgCl_2_ and 0.3% Tween-20 unless otherwise specified. Mg-PBS buffer is composed of DPBS supplemented with 1 mM MgCl_2_.

### *In vitro* transcription IVT

*In vitro* transcription was usually performed in 200 µL transcription mixture containing 4 µg DNA template, transcription buffer (40 mM Tris-HCl pH 8.1, 2 mM spermidine, 22 mM MgCl_2_, 0.01 % Triton X-100), 10 mM DTT, T7 polymerase (0.7 µM home-made stock). After 4 h incubation at 37°C, 2 µL DNase I (10 U/µL) were added into the reaction mixture and incubated for 30 min at 37°C. RNA was purified by gel electrophoresis on a denaturing polyacrylamide gel, excised by UV shadowing and eluted in sodium acetate buffer (0.3 M, pH 5.2) overnight. Finally, RNA was precipitated with isopropanol and then dissolved in water.

### *In vitro* characterization of the dyes and aptamer:dye complexes

For fluorescence measurements, the maximum excitation wavelength was used to excite the dyes or aptamer:dye complexes and their fluorescence intensities were detected at the maximum emission wavelength.

For all *in vitro* experiments, RNA was freshly folded each time. Briefly, RNA was dissolved in water, incubated at 75°C for 2 min and then cooled to 25°C with a cooling rate of 0.1°C/s. Then, 1/5 volume 6× ASBT was added to RNA and incubated for 10 min at 25°C to promote correct RNA folding.

The dissociation constants of aptamer:dye complexes were determined by measuring the fluorescence increase of the dye as function of the RNA concentration. The freshly folded RNA was diluted to the desired concentrations with ASBT and the dye was added to each dilution. The complexes were incubated for 1 h at 25°C before each measurement. The plotted curves were fitted to Hill1 equation for the complexes between the monomeric aptamers and a bivalent dyes, and to equation (1)^1^ for the complexes between dimeric aptamers and bivalent dyes by using least-squared fitting with OriginPro Version 15.

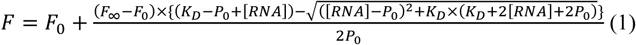

where *F* is the fluorescence at any given aptamer concentration, *F*_0_ is the fluorescence of the free dye with its initial concentration *P*_0_, *F*_∞_ is the maximum fluorescence intensity (full complexation), [RNA] is the final concentration of added aptamer and *K*_D_ is the equilibrium dissociation constant. Used dye concentrations are given in the respective experimental details.

To determine the magnesium ion dependence of the aptamer:dye complexes, RNA was freshly folded in ASBT buffer containing different concentrations of MgCl_2_ as described before. RNA and dye were incubated for 1 h at 25°C before each measurement and were used in equimolar ratios and magnesium chloride concentrations were varied between 0 – 10 mM. The fluorescence intensity of the RNA-dye complex was then measured.

To determine the monovalent cation dependence of the aptamer:dye complexes, RNA was freshly folded in ASBT buffer containing 125 mM MCl (Li^+^, Na^+^, K^+^ or without M^+^) as described before. RNA and the dyes were mixed in equimolar ratios, incubated for 1 h at 25°C, and fluorescence intensities of the RNA:dye complexes were measured.

To determine the temperature dependence of the fluorescence of the aptamer:dye complex, RNA was freshly folded, mixed with the dyes in equimolar ratio, incubated for 1 h at 25°C. The fluorescence intensity was measured while increasing the temperature of the sample from 25°C to 80°C in 30 min.

The fluorescence turn-on values were measured as the ratio of the fluorescence maximum of the fluorophore (20 nM) in the presence (500 nM) and absence of RNA aptamer. The excitation and emission wavelengths of the complex were used.

The D_50_ value was determined as the dielectric constant of water/dioxane mixtures in which **SiR**_**2**_ (2.5 μM) displays half of its maximum absorbance. The absorbance maxima were plotted against the dielectric constants of the respective solvent mixtures containing 100 – 0% dioxane in water (10% v/v increments). The data points were normalized to the maximum absorbance recorded for **SiR**_**2**_.

For binding kinetic studies, the freshly folded aptamer at various concentrations and the dye (200 pM for **1, 2**, and **TMR**_**2**_; 5 nM for **SiR**_**2**_) were rapidly mixed in ASBT at 25°C and the fluorescence intensity at the emission maximum of the complex was recorded over time (every 0.1 s). The observed rate coefficient *k*_obs_ was determined by fitting the curve to equation (2):

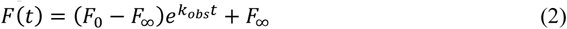

where *F*_0_ is the initial and *F*_∞_ is the final (at equilibrium) fluorescence intensity.

The association rate coefficient *k*_on_ was calculated by fitting the *k*_obs_ values obtained at various RNA concentrations using the equation (3):

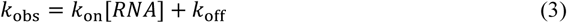

The dissociation rate coefficient *k*_off_ was calculated using the equation (4).

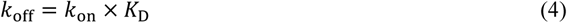

Fluorescence quantum yields were determined after cross-calibrating the set-up for two standards. For TMR-based and SiR-based dyes, sulforhodamine 101 (QY= 1.00 in EtOH)^21^ and oxazine 1 (QY=0.141 in EtOH)^22^ were chosen as the references, respectively. The integral of emission spectra of samples was plotted against their absorbance values at the excitation wavelength (525 nm for TMR-based dyes and 600 nm for SiR-based dyes) at various concentrations (four or more samples with absorbance values of <0.05 to avoid inner filter effects). QYs of the probes were calculated by fitting the resulting plot using the equation (5):

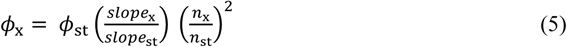

where *Φ* is the fluorescence quantum yield, x and st denote probe and the reference with a known QY, respectively, and *n* is the refractive index of the solvent.

*In vitro* fluorescence intensity decay experiments were conducted as previously described^1^. In brief, freshly folded RNA (500 nM, excess) was mixed with the dyes (20 nM) in ASBT and incubated for 1 h before each measurement. To compare the photostability of biRhoBAST:TMR-dimers and Pepper:HBC620^5^, we continuously illuminated the samples with Luxeon Rebel ES Lime LED (nominal wavelength = 567 nm, U = 3.0 V, I = 800 mA) and the fluorescence intensity of the complexes was measured every 20 min for 3 h.

### DNA Cloning

Double-stranded DNA template carrying biRhoBAST V1 sequence (Supplementary Table 5) flanked by XbaI and Bsp120I restriction sites was generated by PCR, double-digested and ligated into XbaI/Bsp120I double-digested pcDNA3-CFP (Addgene Plasmid #13030) to generate pcDNA3-CFP-biRhoBAST. To construct biRhoBAST repeats, a NheI restriction site was introduced into the downstream of the biRhoBAST sequence in pcDNA3-CFP-biRhoBAST. This new plasmid was then digested with NheI, dephosphorylated and ligated with XbaI/NheI double-digested biRhoBAST V2 sequence (Supplementary Table 5) flanked by XbaI and NheI in order to form pcDNA3-CFP-biRhoBAST_2_. Then, pcDNA3-CFP-biRhoBAST_2_ was double-digested with XbaI and NheI and the biRhoBAST_2_ cassette was gel purified. biRhoBAST_2_ was finally ligated into pcDNA3-CFP-biRhoBAST_2_ which had already been digested with NheI and dephosphorylated to yield pcDNA3-CFP-biRhoBAST_4_. Similarly, the cassette containing biRhoBAST_4_ was obtained by double digestion of pcDNA3-CFP-biRhoBAST_4_ by XbaI and NheI, and subsequent gel purification. It was then ligated into the NheI-digested and dephosphorylated pcDNA3-CFP-biRhoBAST_4_ vector to yield pcDNA3-CFP-biRhoBAST_8_.

The biRhoBAST_*n*_ (*n* = 1, 2, 4 or 8) cassettes were obtained from the double digestion of pcDNA3-CFP-biRhoBAST_*n*_ plasmids with XbaI and NheI, and their subsequent gel purification. To insert biRhoBAST_*n*_ into pAV-U6+27 (Addgene Plasmid #25709), the biRhoBAST cassettes were ligated into XbaI-digested and dephosphorylated pAV U6+27 vector to yield pAV-U6+27-biRhoBAST_*n*_.

Then, pAV-U6+27-biRhoBAST_*n*_ was double-digested with SalI and XbaI, the biRhoBAST_*n*_ cassette was gel purified and ligated into XhoI-digested and dephosphorylated mAzurite-C1 (Addgene Plasmid #54583) to yield mAzurite-biRhoBAST_*n*_. To create mAzurite-24xMS2v6 and mAzurite-biRhoBAST_8_-MS2_24_, a vector containing 24xMS2v6 (Addgene Plasmid #104391) was double-digested and the MS2 cassette was gel purified. It was then ligated into XhoI-digested and dephosphorylated mAzurite or mAzurite-biRhoBAST_8_. For mAzurite-24xMS2v6, a stop codon after the mAzurite coding sequence was introduced via PCR (see Supplementary Table 5 for the primers). To prepare mEGFP-biRhoBAST_*n*_, the respective mAzurite-biRhoBAST_*n*_ vectors were double-digested with NheI and Bsp1407I, and then ligated with NheI/Bsp1407I-digested mEGFP insert that was obtained from a PCR reaction using AAVS1-mEGFP (Addgene Plasmid #91565) as the template DNA.

To yield GFP-β-actin-Zip-biRhoBAST_n_, XbaI/NheI-digested biRhoBAST_*n*_ cassettes were ligated into XbaI-digested and dephosphorylated eTC-GFP-β-actin-Zip (Addgene Plasmid #27123).

For the expression of circular aptamers in mammalian cells, double-stranded DNA templates of biRhoBAST, biSiRA, Mango2 and Pepper (Supplementary Table 5) were generated via PCR, digested with SacII and NotI and ligated into SacII/NotI-digested pAV-U6+27-Tornado-Broccoli (Addgene Plasmid #124360) to yield pAV-U6+27-Tornado-biRhoBAST, pAV-U6+27-Tornado-biSiRA, pAV-U6+27-Tornado-Mango2 and pAV-U6+27-Tornado-Pepper, respectively.

### Confocal imaging of live cells

HEK293T, U2OS, Cos7 and MEF cells were cultured at 37°C with 5% CO_2_ in Dulbecco’s Modified Eagle’s Medium (high glucose), HEPES and glutamine without phenol red (Thermo Fisher Scientific) supplemented with 10% FBS (Gibco), 50 U ml^−1^ of penicillin and 50 μg ml^−1^ of streptomycin (Thermo Fisher Scientific). Cells were routinely passaged after 2-3 days or after reaching ∼80 % confluency using TrypLE Express (Gibco) for the detachment of cells from culture flasks. Transfection was carried out with FuGENE HD (Promega) according to the manufacturer’s instructions.

For live-cell imaging experiments, 5×10^4^ (for 8-well chamber) or 2×10^4^ (for 18-well chamber) cells were seeded into µ-Slide Glass Bottom from IBIDI. Before seeding the HEK293T cells, the glass surfaces were treated with KOH (10 M) for 10 min, thoroughly washed with water and coated with poly-D-lysine. One day after seeding, cells were transfected using FugeneHD (Promega) according to the manufacturer’s instructions. After 24-48 h, the cells were washed twice with Leibowitz medium (L15) and the medium was exchanged to L15 medium supplemented with the nuclear stain (1 µg ml^−1^ of Hoechst 33342, 1 µM SYTO9 or 5 µM Draq5) and the appropriate concentrations of the desired dyes (indicated in the figure captions of the respective experiments). The cells were then incubated for 1-2 h in a humidified 37°C chamber without CO_2_ and then imaged at 37°C in a live-cell imaging chamber with controlled humidity either with a point-scanning confocal or spinning disc confocal microscope.

For point-scanning, a Nikon A1R mounted on a fully automated inverted Nikon Ti2 was used with a hybrid scanner (galvano/resonant) equipped with a 60x oil objective (Nikon N Apo ×60 NA 1.4 λs OI) and controlled by NIS software (Nikon). To excite Hoechst 33342 or mAzurite, we used a 405 nm laser with an emission filter set of 450 ± 25 nm; to excite GFP or SYTO9, we used a 488 nm laser with an emission filter set of 525 ± 25 nm; to excite TMR or mScarlet, we used a 561 nm laser with an emission filter set of 595 ± 25 nm; to excite SiR or Draq5, we used a 640 nm laser with an emission filter set of 700 ± 37.5 nm. Z-Stack images were taken with steps sizes of 200-500 nm and images were analyzed with Fiji/ImageJ^23^. Presented images were unprocessed as average projections of 5-7 slices with 500 nm step size for HEK293T and 200 nm step size for Cos7. For quantifications, the background was corrected by subtracting the fluorescence intensity of control cells which had not been incubated with the dyes from the whole image.

### smFISH imaging

Single-molecule FISH probes (HuluFISH) for GFP were designed and enzymatically labelled with Atto647 by PixelBiotech (see Supplementary Table 5for sequences). MEF cells were transfected with the appropriate plasmid containing GFP-β-actin-biRhoBAST_8_ as described above. Cells were fixed with 4% PFA 36 h after transfection and washed twice with Mg-PBS containing 100 mM glycine. The fixed cells were permeabilized with 300 µL of 0.1% Triton X-100 in Mg-PBS for 15 min at room temperature and subsequently washed three times with 500 µL Mg-PBS. Then, the fixed and permeabilized cells were incubated with HuluWash for 10 min. After removing HuluWash, 120 µL HuluHyb containing smFISH probes was added into each well and incubated for 16 h at 30°C in a humidified chamber. Then, the cells were washed three times with HuluWash and incubated with Mg-PBS for 30 min at 30°C. Before imaging, cells were incubated with 1 µg/mL Hoechst 33342 and 50 nM **TMR**_**2**_ in Mg-PBS for 30 min at room temperature. Images were taken with the scanning confocal microscope described above with 60x oil objective, 0.05 µm per pixel and 17 z-stacks with 0.2 µm step size.

Images were background-subtracted with rolling ball (15 pixel radius). Colocalization test was performed by JACoP ImageJ plugin^24^.

### Single mRNA imaging

For fast image acquisition, a spinning disc confocal microscope was used. A PerkinElmer ERS-VoX attached to an automated inverted Nikon Ti microscope equipped with a Nikon Plan Apo VC 100x NA 1.4 oil immersion objective and a Hamamatsu C9100-02 EMCCD camera and a Yokagawa CSU-X1 confocal scanning unit was controlled by PerkinElmer Volocity software. Temperature and humidity control were provided by a TokaiHit on stage incubation system. Lasers of 405, 488, 561 and 640 nm wavelength were used for the abovementioned dyes in combination with a blue/red dual pass filter (445 ± 30 nm and 615 ± 35 nm), a green filter (527 ± 27.5 nm) and a cyan/red dual pass filter (485 ± 30 nm and 705 ± 45 nm). To detect GFP and TMR colocalization in live cells, a second Hamamatsu C9100-02 EMCCD camera was used which allowed simultaneous imaging of both green and red colors. Unless noted otherwise, single z-plane images were acquired typically for 15-60 s with exposure times of 100 ms (9.7 frames per second). For live cell single particle tracking, we used TrackMate ImageJ plugin^25^. A LoG filter was applied to detect single particles. Foci diameter 500 nm, frame rates (0.103 s per frame) were automatically included from metadata. For building tracks, foci were given a maximum of 1 µm for linking frame-to-frame displacement and 1 µm gap-closing. To recover partial loss of trajectories a maximum frame gap of 5 frames was allowed. The obtained trajectories were filtered eventually to show at least 10 localizations as well as immobile or low-quality foci were excluded. The trajectories were then analyzed with MSDanalyzer in Matlab^26^. Similar to smFISH, for dual-color tracking a colocalization test was performed with JACoP ImageJ plugin^24^.

### Photostability of FLAPs in live cells

A modified procedure of Braselmann et al.^4^ was carried out to compare the photostability of various FLAPs in live cells. In brief, the spinning disc confocal microscope described above was used with a Nikon Plan Apo λ 60x NA 1.40 oil immersion objective to record fluorescence decay curves in live HEK293T cells. Cells were transfected with Tornado-Broccoli, Tornado-Mango2, Tornado-Pepper, Tornado-biRhoBAST or Tornado-biSiRA and incubate with BI (10 µM), TO1-B (200 nM), HBC620 (1.0 µM nM), TMR_2_ (50 nM) or SiR_2_ (100 nM), respectively. We also used mEGFP or mScarlet (Addgene Plasmid #85042) plasmids as a transfection control to discriminate the transfected cells from the untransfected ones. Before the experiment, transfected cells were only visualized in Hoechst and the co-transfected fluorescent proteins with low laser powers to avoid prebleaching of the FLAPs. The laser power for each experiment was adjusted such that each FLAP has the same initial excitation rate. In a second experiment, the laser power was kept constant (1.65 mW) for all FLAPs. Cells were imaged for 90 s with an exposure time of 100 ms (9.7 frames per second). For data evaluation, the average fluorescence intensities in the cytosol of more than 50 cells were determined for every frame and the fluorescence decay over time was evaluated. Average initial fluorescence of each FLAP was normalized to 1.

## References

1. Sunbul, M. et al. Super-resolution RNA imaging using a rhodamine-binding aptamer with fast exchange kinetics. Nat. Biotechnol. 39, 686–690 (2021).

2. Strack, R.L., Disney, M.D. & Jaffrey, S.R. A superfolding Spinach2 reveals the dynamic nature of trinucleotide repeat–containing RNA. Nat. Methods 10, 1219–1224 (2013).

3. Song, W. et al. Imaging RNA polymerase III transcription using a photostable RNA–fluorophore complex. Nat. Chem. Biol. 13, 1187–1194 (2017).

4. Braselmann, E. et al. A multicolor riboswitch-based platform for imaging of RNA in live mammalian cells. Nat. Chem. Biol. 14, 964–971 (2018).

5. Chen, X. et al. Visualizing RNA dynamics in live cells with bright and stable fluorescent RNAs. Nat. Biotechnol. 37, 1287–1293 (2019).

6. Wu, J. et al. Live imaging of mRNA using RNA-stabilized fluorogenic proteins. Nat. Methods 16, 862–865 (2019).

7. Wirth, R. et al. SiRA: a silicon rhodamine-binding aptamer for live-cell super-resolution RNA imaging. J. Am. Chem. Soc. 141, 7562–7571 (2019).

8. Terdale, S. & Tantray, A. Spectroscopic study of the dimerization of rhodamine 6G in water and different organic solvents. J. Mol. Liq. 225, 662–671 (2017).

9. Bouhedda, F. et al. A dimerization-based fluorogenic dye-aptamer module for RNA imaging in live cells. Nat. Chem. Biol. 16, 69–76 (2020).

10. Setiawan, D., Kazaryan, A., Martoprawiro, M.A. & Filatov, M. A first principles study of fluorescence quenching in rhodamine B dimers: how can quenching occur in dimeric species? Phys. Chem. Chem. Phys. 12, 11238–11244 (2010).

11. Kitov, P.I. & Bundle, D.R. On the nature of the multivalency effect: a thermodynamic model. J. Am. Chem. Soc. 125, 16271–16284 (2003).

12. Mack, E.T. et al. Dependence of avidity on linker length for a bivalent ligand–bivalent receptor model system. J. Am. Chem. Soc. 134, 333–345 (2012).

13. Litke, J.L. & Jaffrey, S.R. Highly efficient expression of circular RNA aptamers in cells using autocatalytic transcripts. Nat. Biotechnol. 37, 667–675 (2019).

14. Li, X. et al. Fluorophore-Promoted RNA Folding and Photostability Enables Imaging of Single Broccoli-Tagged mRNAs in Live Mammalian Cells. Angew. Chem. Int. Ed. 59, 4511–4518 (2020).

15. Cawte, A.D., Unrau, P.J. & Rueda, D.S. Live cell imaging of single RNA molecules with fluorogenic Mango II arrays. Nat. Commun. 11, 1–11 (2020).

16. Lawrence, J.B. & Singer, R.H. Intracellular localization of messenger RNAs for cytoskeletal proteins. Cell 45, 407–415 (1986).

17. Rodriguez, A.J., Shenoy, S.M., Singer, R.H. & Condeelis, J. Visualization of mRNA translation in living cells. J. Cell Biol. 175, 67–76 (2006).

18. Tutucci, E. et al. An improved MS2 system for accurate reporting of the mRNA life cycle. Nat. Methods 15, 81–89 (2018).

19. Tutucci, E., Vera, M. & Singer, R.H. Single-mRNA detection in living living. cerevisiae using a re-engineered MS2 system. Nat. Protoc. 13, 2268–2296 (2018).

20. Fusco, D. et al. Single mRNA molecules demonstrate probabilistic movement in living mammalian cells. Curr. Biol. 13, 161–167 (2003).

## Additional References

21. Karstens, T. & Kobs, K.J. Rhodamine B and rhodamine 101 as reference substances for fluorescence quantum yield measurements. J. Phys. Chem. 84, 1871–1872 (1980).

22. Rurack, K. & Spieles, M. Fluorescence quantum yields of a series of red and near-infrared dyes emitting at 600− 1000 nm. Anal. Chem. 83, 1232–1242 (2011).

23. Schindelin, J. et al. Fiji: an open-source platform for biological-image analysis. Nat. Methods 9, 676–682 (2012).

24. Bolte, S. & Cordelières, F.P. A guided tour into subcellular colocalization analysis in light microscopy. J. Microsc. 224, 213–232 (2006).

25. Tinevez, J.-Y. et al. TrackMate: An open and extensible platform for single-particle tracking. Methods 115, 80–90 (2017).

26. Tarantino, N. et al. TNF and IL-1 exhibit distinct ubiquitin requirements for inducing NEMO–IKK supramolecular structures. J. Cell Biol. 204, 231–245 (2014).

