## Supplementary Information for "Bright, fluorogenic and photostable avidity probes for RNA imaging"

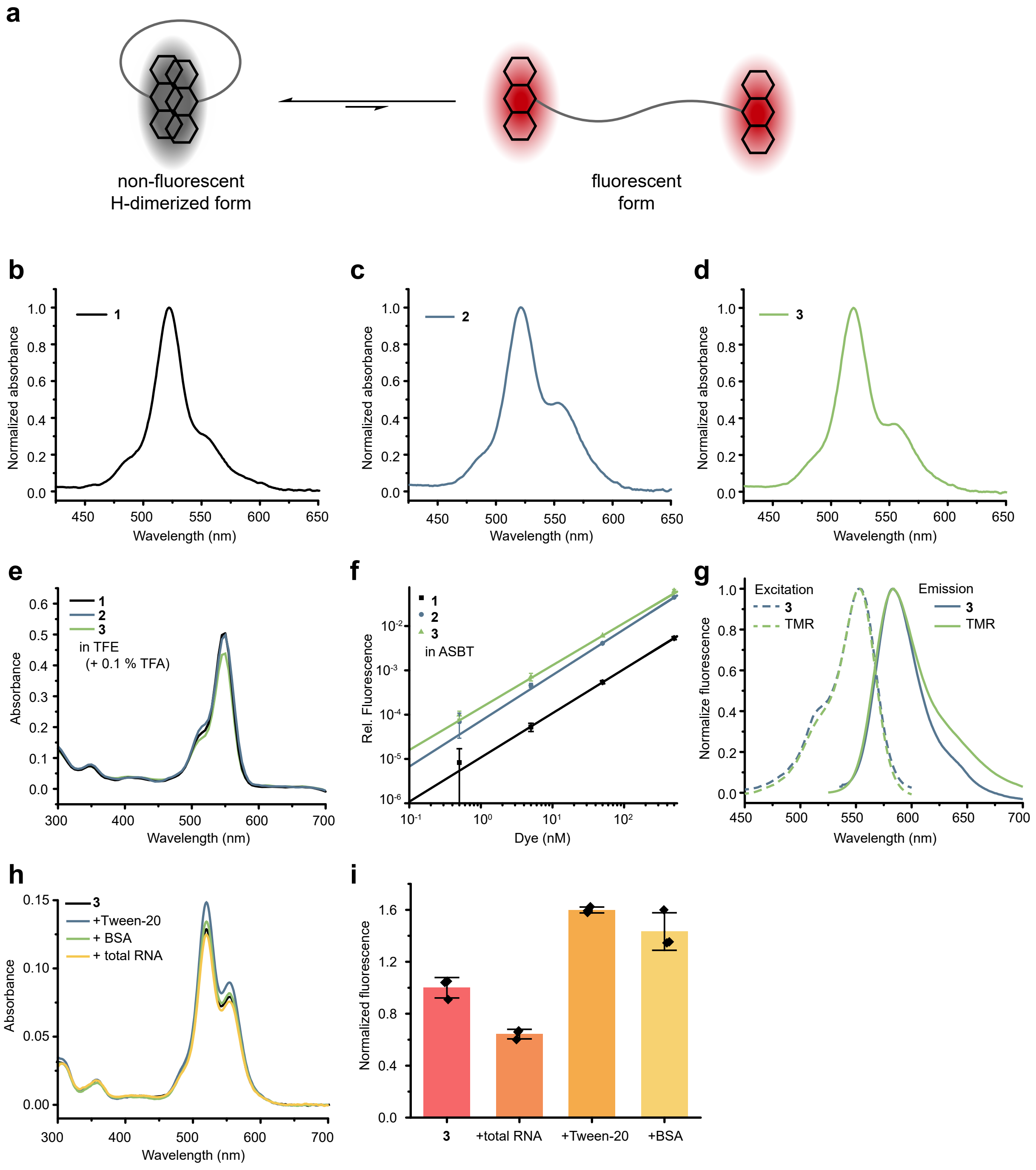


**Supplementary Figure 1:** Photophysical characterization of H-dimer forming bivalent dyes **1**, **2** and **3**.

**a)** Illustration showing an equilibrium between a non-fluorescent H-dimer form and a fluorescent form of TMR dimers. Two TMR molecules were connected via a short polyethylene glycol linker. Therefore, TMR dimers predominantly exist in the H-dimer form due to high local concentrations of the rhodamine dyes. **b-d)** Absorption spectra of TMR-dimers **1**, **2** and **3** (1 µM) in aqueous solution, showing absorption maxima at 517 nm. The blue-shifted absorption maximum compared to monomeric TMR (see Figure 1d) indicates successful H-dimer formation[1, 2]. **e)** Absorption spectra of TMR dimers **1**, **2** and **3** (2 µM) in trifluoroethanol supplemented with 0.1% TFA to prevent H-dimer formation[1, 3]. Absorption maxima around 554 nm indicate the disruption of H-dimer form. **f)** Comparison of fluorescence intensities of TMR-dimers **1**, **2** and **3** at different concentrations. Shorter linkers result in the lower residual fluorescence: **1** < **2** < **3**. Fluorescence values are given relative to the fluorescence of 1 µM monomeric TMR to display the low fluorescence of the quenched dyes. In addition, the linear correlation between the fluorescence and the concentration indicated intramolecular H-dimer formation[1]. In contrast, intermolecular H-aggregates are typically present above micromolar concentrations and their fluorescence is affected by dilution, displaying positive deviations from linearity in concentration versus fluorescence intensity plots[3]. Data represent the mean ± s.d. of three independent measurements. **g)** Excitation and emission spectra of **3** ($\lambda_{max}^{exc}$= 553 nm and $\lambda_{max}^{em}$ = 583 nm) in aqueous solution are similar to those of TMR. This suggests that the low background fluorescence of **3** originates from incomplete H-dimer formation, not from J-dimers, since J-dimers would display red-shifted excitation maxima compared to free TMR[1]. The effect of total RNA (500 ng μL$^{-1}$), BSA (1.0 μg μL$^{-1}$) and Tween-20 (0.1%) on **h)** the absorption spectra, and **i)** the fluorescence intensity of **3** (600 nM). These additives did not cause a dramatic change in neither absorption spectrum nor the fluorescence intensity of **3**, emphasizing the stability of intramolecular TMR H-dimers. Data represent the mean ± s.d. of three independent measurements.

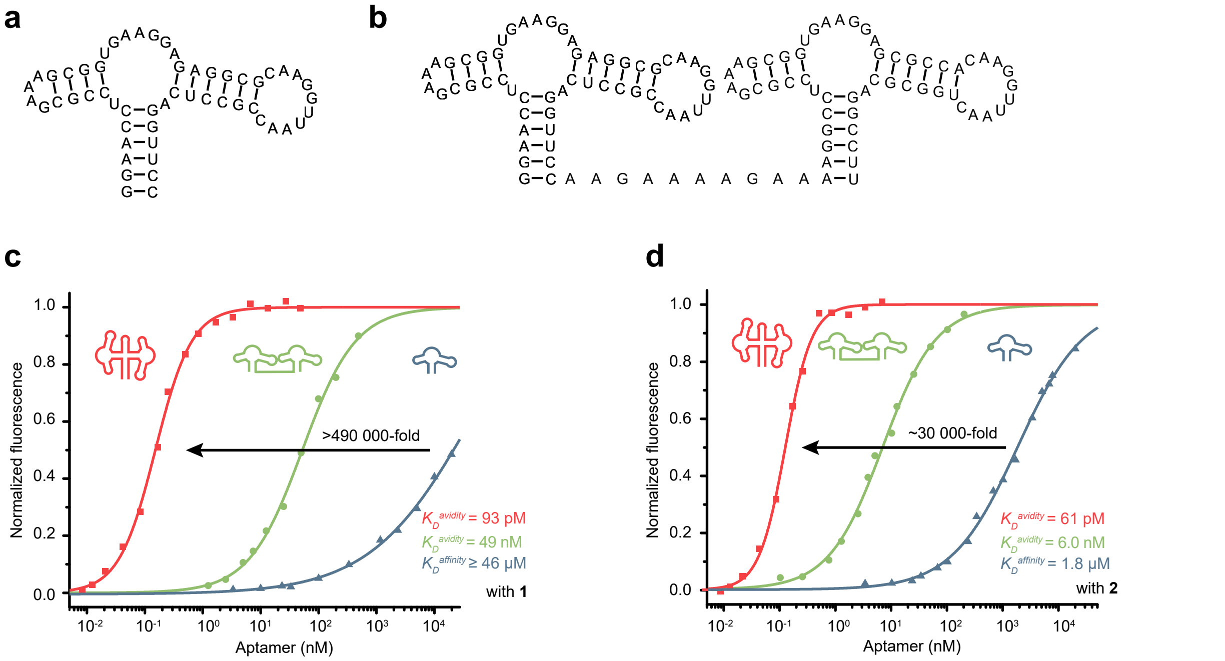


**Supplementary Figure 2:** Avidity dramatically enhances the binding strength between the bivalent TMR dyes (**1** and **2**) and dimeric RhoBAST aptamers.

Predicted secondary structures of **a)** RhoBAST; **b)** its dimeric variant with a 10-nucleotide linear and flexible linker. Dissociation constant ($K_D$) measurements between RhoBAST variants (monomeric in blue, dimeric with a flexible linker in green, and biRhoBAST with a rigid linker in red) and **c)** bivalent TMR dye **1**; **d)** bivalent TMR dye **2**. Plots were obtained by titrating dye solutions with increasing concentrations of aptamers and recording the fluorescence intensities. The final concentrations of the dyes (**1** and **2**) during measurements were 5 nM, 1 nM and 0.1 nM when RhoBAST, dimeric RhoBAST with a flexible linker and biRhoBAST were used as aptamers, respectively. $K_D^{avidity}$ depends significantly on the rigidity, geometry and preorganization of the aptamers as well as on the linker length of the dyes, which also determines the strength of the H-dimers[4, 5].

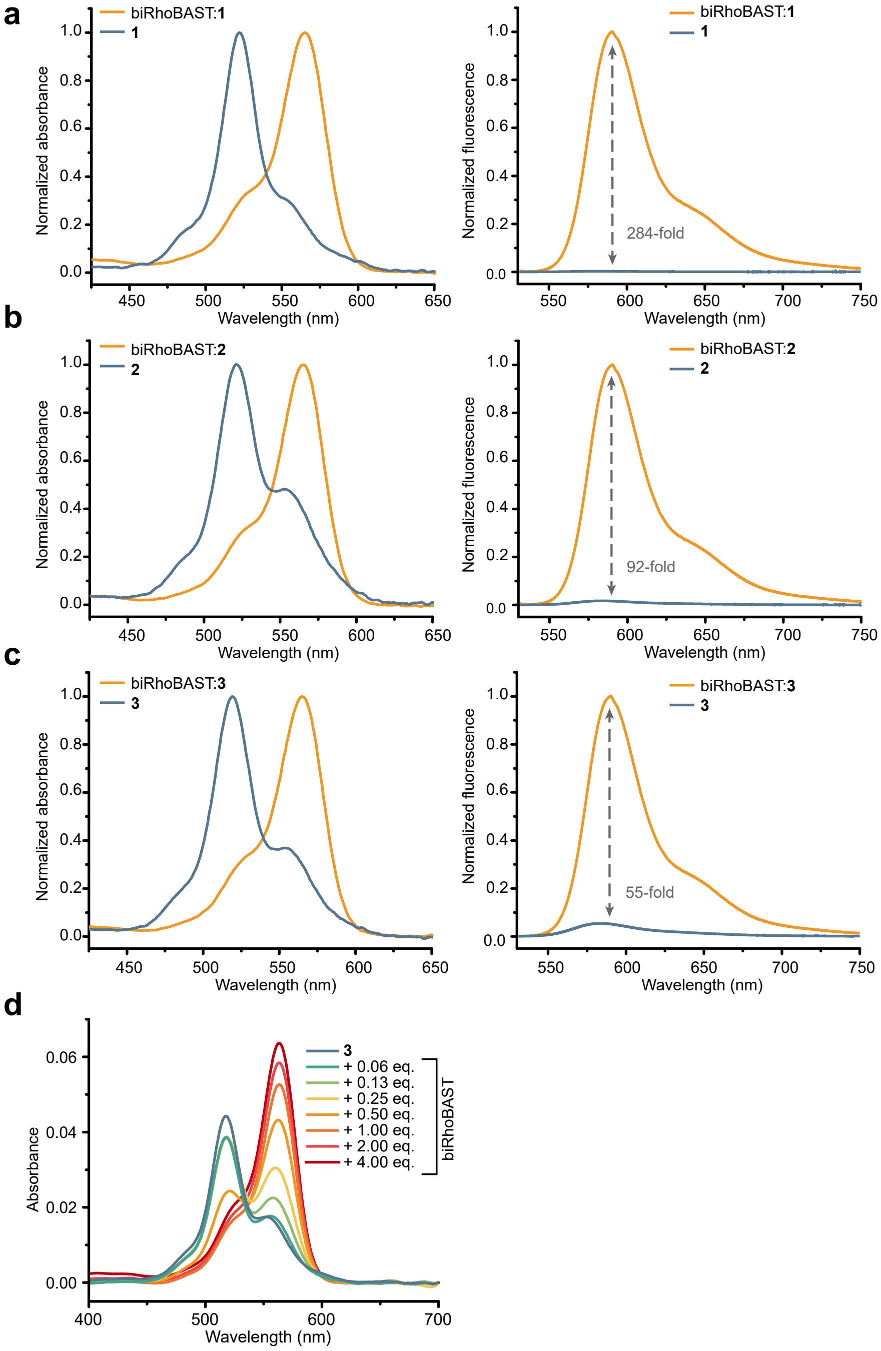


**Supplementary Figure 3:** The fluorescence of bivalent TMR dyes **1**, **2** and **3** increases upon binding to biRhoBAST.

**a-c)** Absorption and emission spectra of bivalent TMR-dimers **1**, **2**, and **3** (1 µM) and their complexes with biRhoBAST (5 µM). The binding of biRhoBAST to the dyes effectively destroys the H-dimers, resulting in fluorescence enhancement (shown by grey arrows). **d)** Titration of **3** (400 nM) with increasing amounts of biRhoBAST indicating the ability of biRhoBAST to break TMR H-dimers. The disruption of the H-dimer was confirmed by the increase of the complex absorption at 564 nm and the decrease in the H-dimer band at 517 nm.

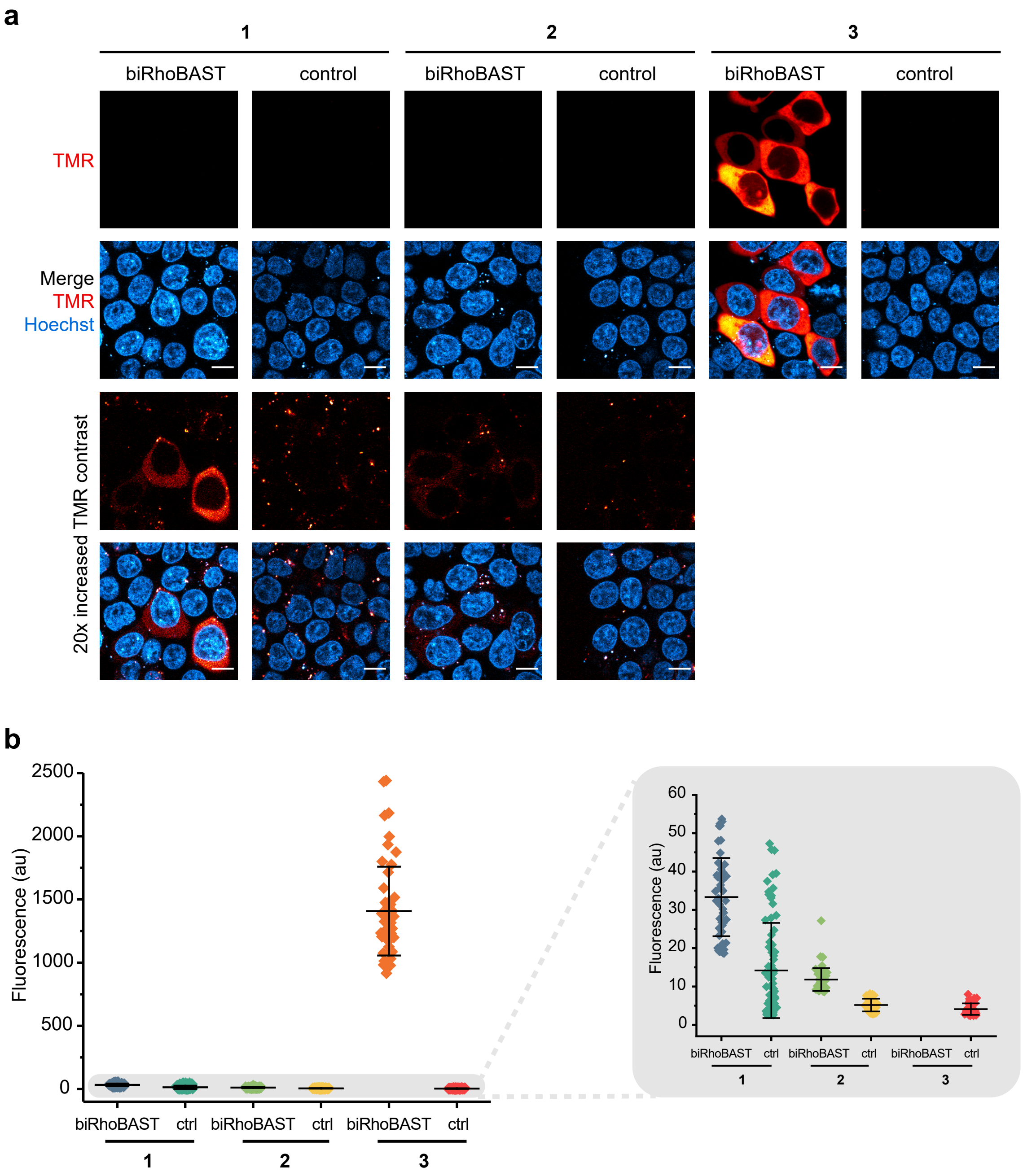


**Supplementary Figure 4:** Bivalent TMR dye **3** (**TMR$_2$**) is superior to **1** and **2** for RNA imaging in live cells.

**a)** HEK293T cells expressing circular biRhoBAST or Broccoli (control)[6] were incubated with 50 nM of **1**, **2** or **3** (**TMR$_2$**) and imaged using a point-scanning confocal microscope. While **TMR$_2$** enabled imaging biRhoBAST with excellent signal-to-background ratios in live cells, dyes **1** and **2** did not yield satisfactory signal-to-background ratios. Transfected cells were found by co-transfected mEGFP. Scale bars, 10 µm. **b)** The fluorescence intensity (mean ± s.d) in cells ($n \geq 50$ cells) described in panel **a**. Grey box shows a zoom-in of the plot.

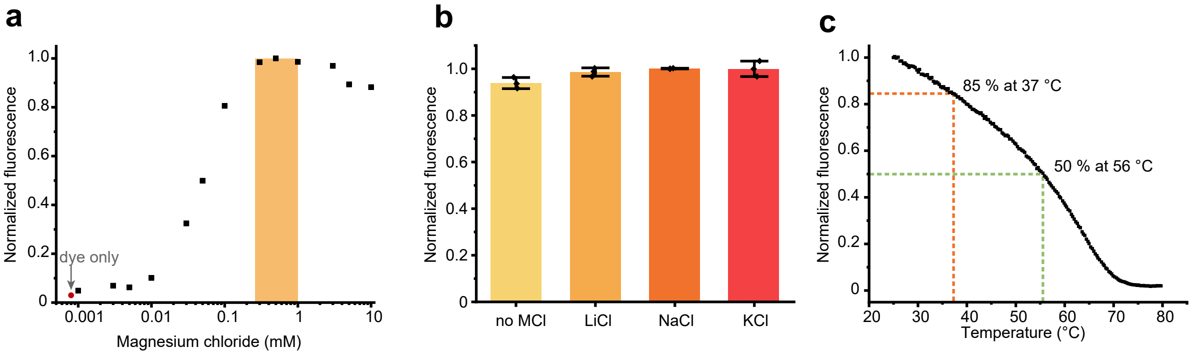


**Supplementary Figure 5:** Characterization of the biRhoBAST:**TMR₂** complex.

**a)** Dependence of the biRhoBAST:**TMR₂** (100 nM) fluorescence intensity on magnesium ion concentration. The physiologically relevant magnesium concentration range in mammalian cells (0.25 – 1 mM) is highlighted in orange[7]. **b)** Dependence of the biRhoBAST:**TMR₂** (100 nM) fluorescence intensity on monovalent cations $Li^+$, $Na^+$ and $K^+$. biRhoBAST:**TMR₂** has no preference for a monovalent cation and showed high fluorescence even in the absence of monovalent cations, suggesting that biRhoBAST does not fold into a G-quadruplex structure[8]. Data represent the mean ± s.d. of three independent measurements. **c)** Dependence of the biRhoBAST:**TMR₂** (100 nM) fluorescence intensity on temperature. The initial fluorescence intensity of the complex decreased by only 15% at 37 °C and 50% at 56 °C.

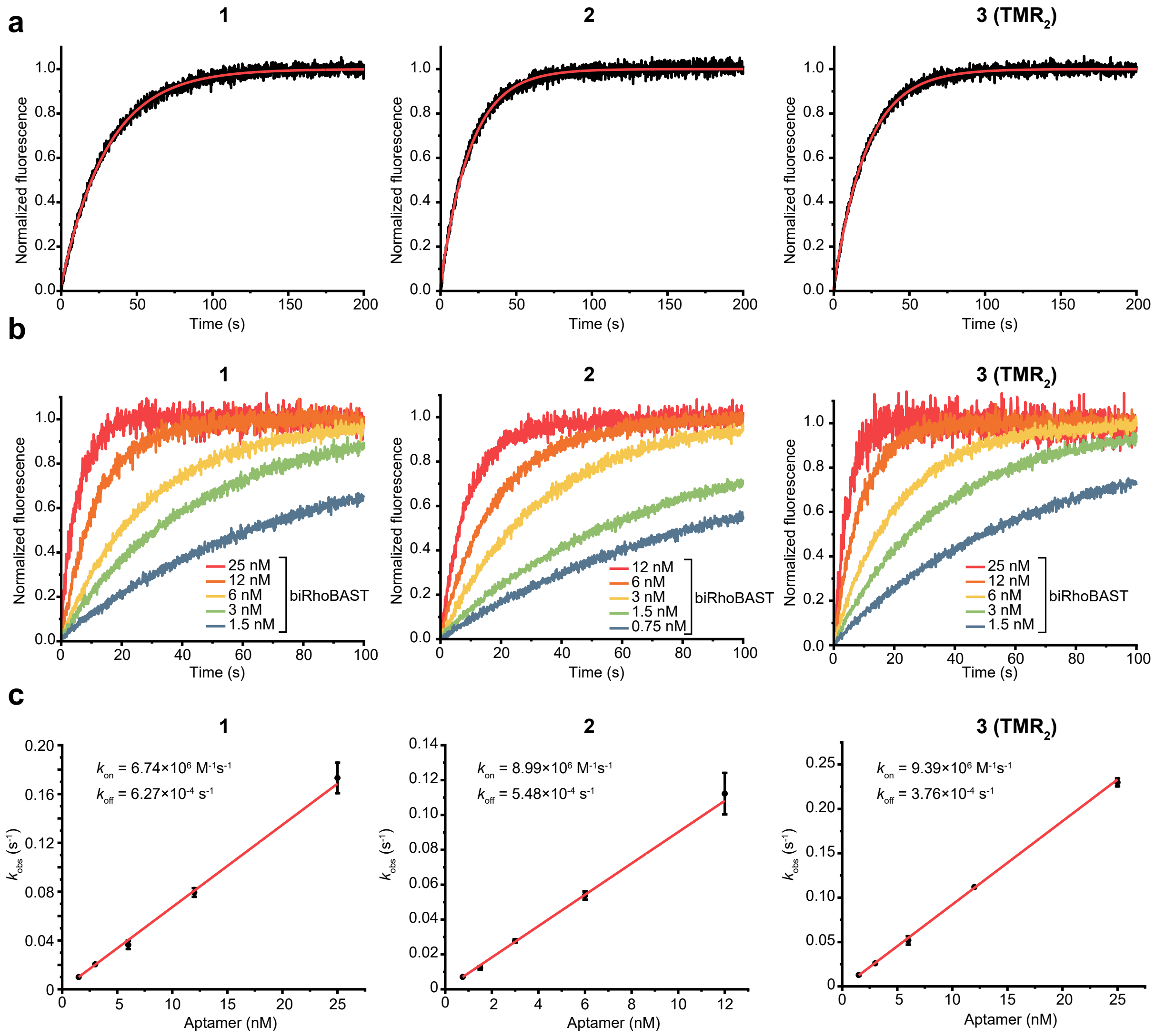


**Supplementary Figure 6:** Determination of association and dissociation rate coefficients for biRhoBAST:**1**, biRhoBAST:**2** and biRhoBAST:**3.**

**a)** Fluorescence increase upon mixing biRhoBAST (6 nM) and the respective bivalent TMR dye (200 pM) due to complex formation over time. **b)** Fluorescence increase upon mixing bivalent TMR dyes (200 pM) with different concentrations of biRhoBAST over time. **c)** Observed rate coefficients ($k_{obs}$) were obtained by fitting the curves in panel **b** to A (1 – (exp [-$k_{obs}$ t]) where A is the amplitude at t = ∞. These $k_{obs}$ values were then plotted against the respective RNA concentrations and the $k_{on}$ values were calculated from the slope of the linear fit; whereas $k_{off}$ values were calculated from the equation $k_{off} = k_{on} K_D$. Data represent mean ± s.d. from three independent measurements.

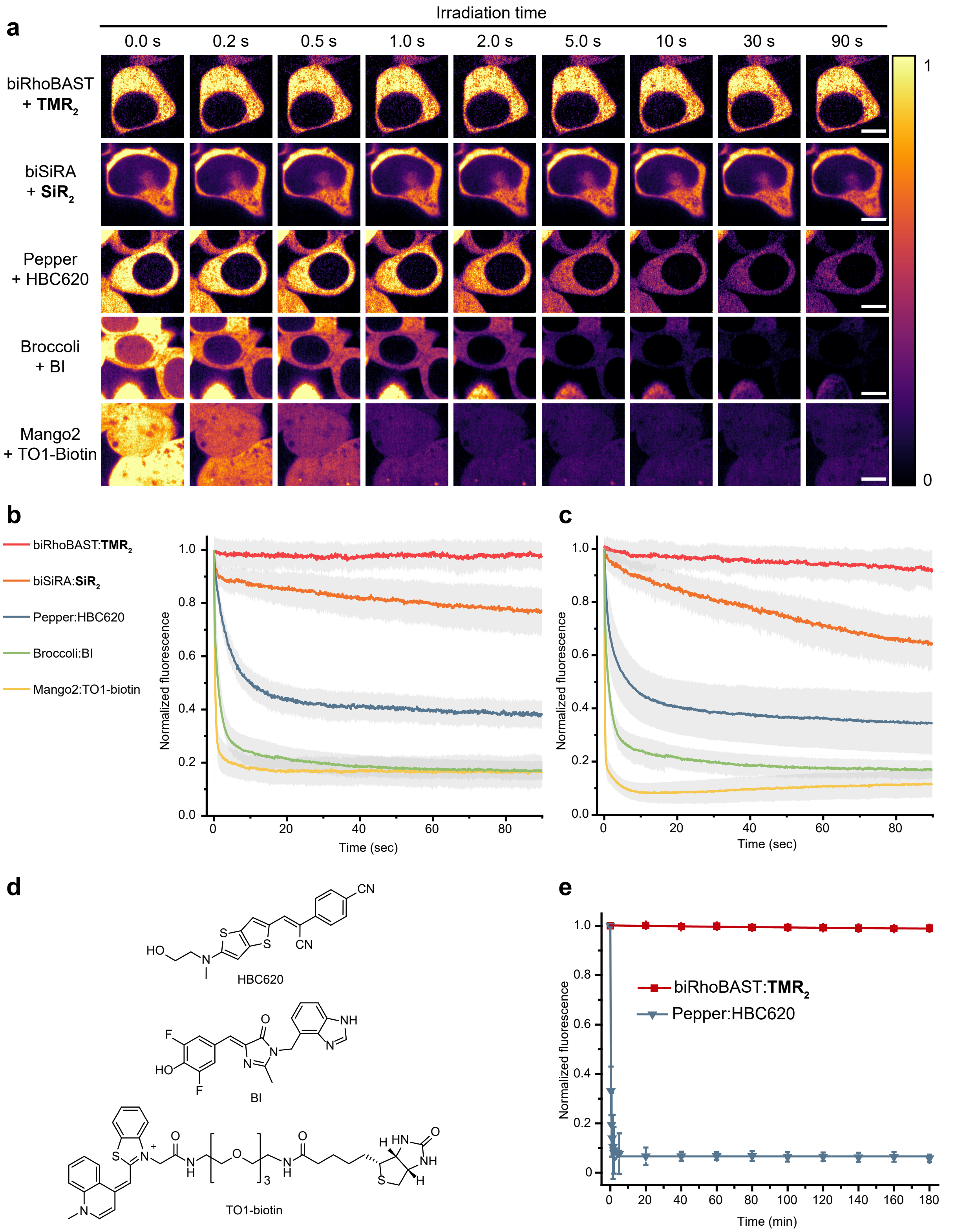


**Supplementary Figure 7:** biRhoBAST:**TMR₂** and biSiRA:**SiR₂** are remarkably photostable in live cells and *in vitro*.

**a)** HEK293T cells expressing the circular biRhoBAST, biSiRA, Pepper, Broccoli or Mango2 were incubated with their cognate dyes **TMR₂** (100 nM), **SiR₂** (200 nM), HBC620 (1.0 µM)[9], BI (10 µM)[10] or TO1-biotin (200 nM)[11], respectively, and imaged continuously for 90 s with 9.7 frames/second on a spinning dis k confocal microscope. The laser power for each image was adjusted such that each aptamer:dye complex was excited with the same initial rate[12]. Transfected cells were found by co-transfected mEGFP for biRhoBAST:**TMR₂**, biSiRA:**SiR₂** and

Pepper:HBC620, and mScarlet for Broccoli:BI and Mango2:TO1-biotin. **b)** Fluorescence intensity (mean ± s.d.) decay curves of the acquired data in panel **a** (*N* ≥ 50 cells). The average fluorescence intensities in cells at t = 0 were normalized to 1. **c)** Fluorescence intensity (mean ± s.d.) decay curves of FLAPs in live cells when they all were illuminated with the same laser power (1.65mW). **d)** Chemical structures of the used dyes. **e)** Fluorescence intensity decay curves of biRhoBAST:$\mathbf{TMR_2}$ and Pepper:HBC620[9] upon irradiation with a 561 nm LED *in vitro*. Dyes (20 nM) were incubated with excess aptamers (500 nM) and the fluorescence intensity was measured after constant irradiation with a 561 nm LED. Data represent the mean ± s.d. of three independent measurements.

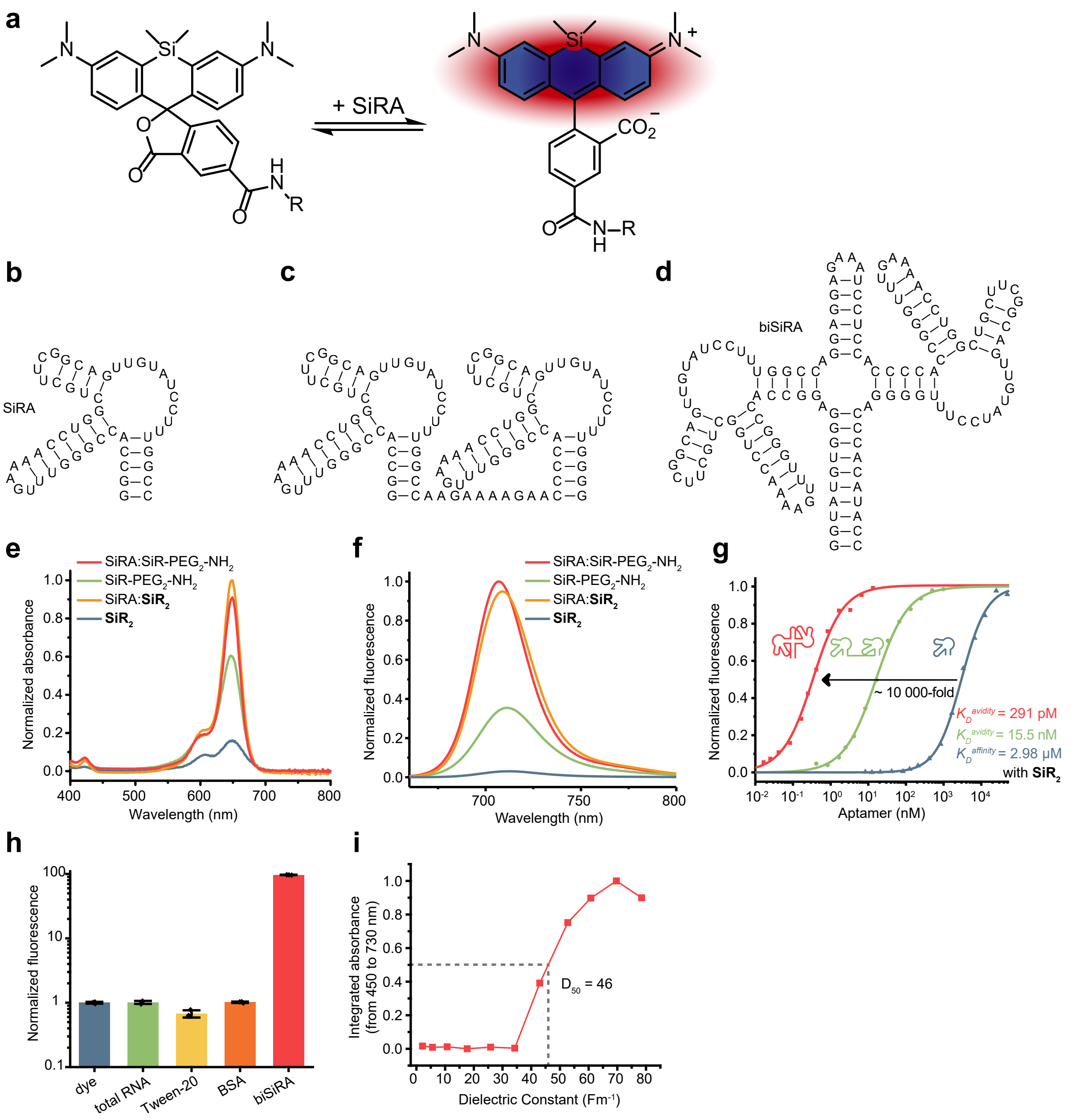

**Supplementary Figure 8:** Characterization of the highly fluorogenic **SiR$_2$** dye.

**a)** Fluorogenicity of silicone rhodamines results from spirolactonization, which was shown to be influenced by SiRA[13]. Predicted secondary structure of **b)** SiRA, **c)** its dimeric variant with a 10-nucleotide linear and flexible linker and **d)** its dimeric variant biSiRA consisting of a four-way junction with a stabilizing stem loop that locks the orientation and distance between two SiRA units. **e)** Absorption and **f)** emission spectra of dimeric SiR (**SiR$_2$,** 1 µM) in comparison to monomeric SiR-PEG$_2$-NH$_2$ (2 µM) in the presence and absence of SiRA (50 µM), demonstrating the ability of SiRA to enhance the absorbance and fluorescence of the dyes. **SiR$_2$** showed a higher absorbance and fluorescence turn-on, most likely a product of spirolactonization and H-dimer formation (see Supplementary Note 1). **g)** Dissociation constant ($K_D$) measurements of complexes between SiRA variants (monomeric in blue, dimeric with a flexible linker in green, and biSiRA with a rigid linker in red) and the bivalent **SiR$_2$** dye. The plot was obtained by titrating different dye solutions with increasing concentrations of aptamers and recording the fluorescence intensities. The final concentrations of the dyes (**1** and **2**) during measurements were 50 nM, 1 nM and 0.1 nM when SiRA, dimeric SiRA with a flexible linker and biSiRA were used as aptamers, respectively. The $K_D^{avidity}$ significantly depends on the rigidity, geometry and preorganization of the aptamers[4, 5]. **h)** The fluorescence intensity of **SiR$_2$** (60 nM) did not change in the presence of total RNA (42 ng µL$^{-1}$), BSA (17 µg µL$^{-1}$) or Tween-20 (0.1%), but increases 100-fold in presence of biSiRA (300 nM, 14 ng µL$^{-1}$). Data represent the mean ± s.d. of three independent measurements. **i)** Absorption spectra of the colored form of **$SiR_2$** (2.5 µM) in water–dioxane mixtures (v/v, 0/100–100/0; in 10% steps) as a function of the dielectric constant. The absorption spectra were integrated from 450 to 730 nm, normalized to the maximum integrated absorption and plotted against the dielectric constant of the respective solvent mixture.

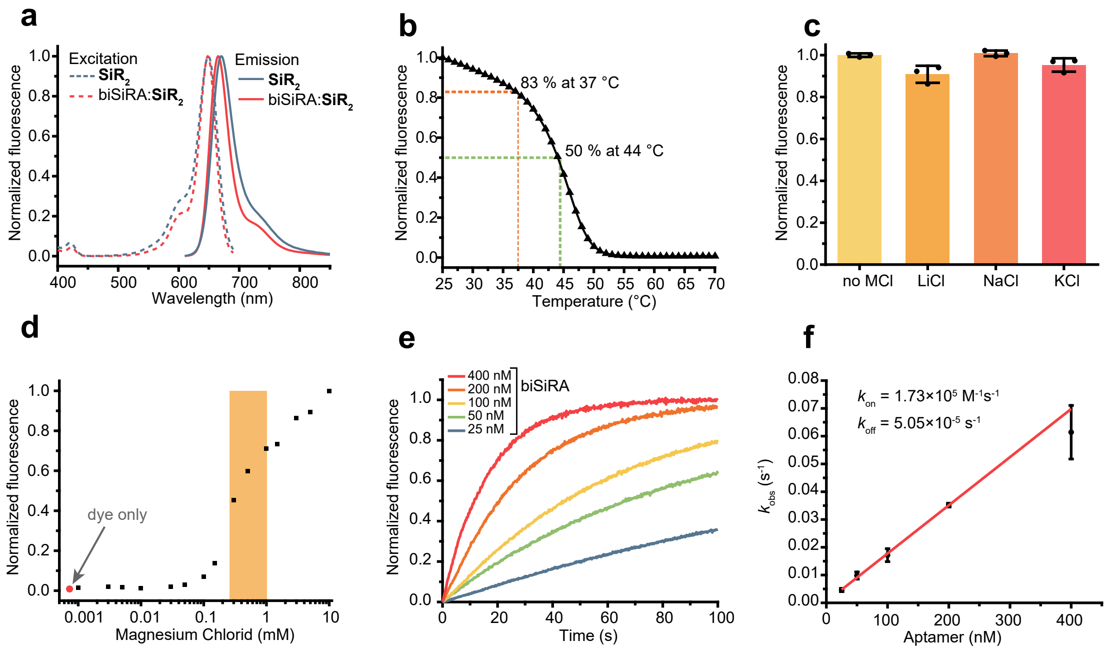


**Supplementary Figure 9:** Characterization of the biSiRA:**SiR₂** complex.

**a)** Excitation and emission spectra of **SiR₂** (600nM) and the biSiRA:**SiR₂** (600 nM) complex, prepared by mixing 600 nM **SiR₂** and 3 µM SiRA. **b)** Dependence of the biSiRA:**SiR₂** (100 nM) fluorescence intensity on temperature. The initial fluorescence intensity of the complex decreased by only 17% at 37 °C and 50% at 44 °C. **c)** Dependence of biSiRA:**SiR₂** (100 nM) fluorescence intensity on monovalent cations $Li^+$, $Na^+$ and $K^+$. biSiRA:**SiR₂** has no preference for a monovalent cation and showed high fluorescence even in the absence of monovalent cations, suggesting that biSiRA does not fold into a G-quadruplex structure[8]. Data represent the mean ± s.d. of three independent measurements. **d)** Dependence of biSiRA:**SiR₂** (100 nM) fluorescence intensity on magnesium ion concentration. The physiologically relevant magnesium concentration range in mammalian cells (0.25 – 1 mM) is highlighted in orange[7]. **e)** Fluorescence increase upon mixing 5 nM **SiR₂** with different concentrations of biSiRA over time. **f)** Observed rate coefficients ($k_{obs}$) were obtained by fitting the curves in panel **e** to A (1 – (exp [-$k_{obs}$ t]) where A is the amplitude at t = ∞. These $k_{obs}$ values were then plotted against the respective RNA concentrations and the $k_{on}$ values were calculated from the slope of the linear fit; whereas $k_{off}$ values were calculated from the equation $k_{off}$ = $k_{on}$ $K_D$. Data points represent mean ± s.d. from three independent measurements.

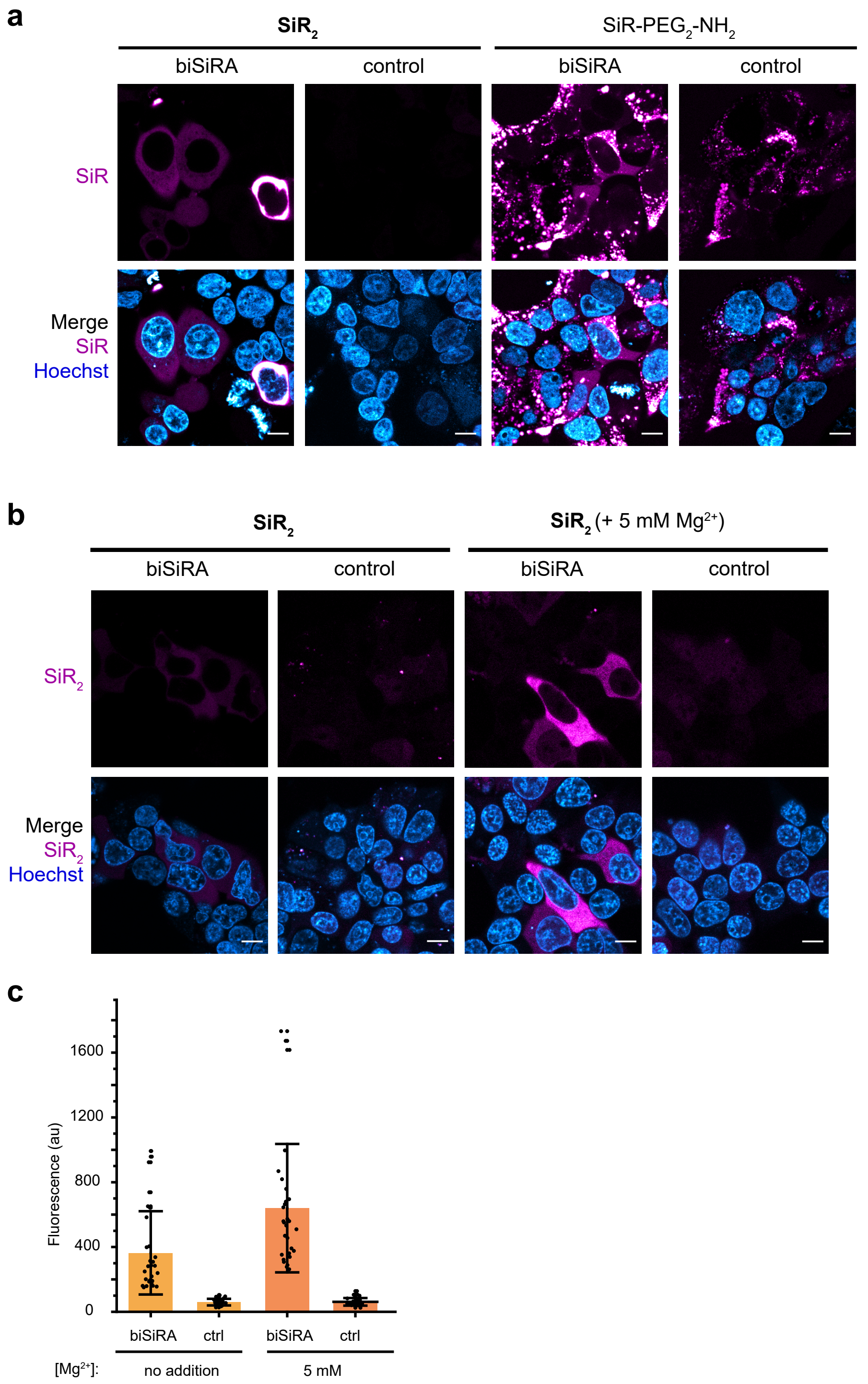


**Supplementary Figure 10:** biSiRA:**SiR₂** enables RNA imaging in live mammalian cells.

**a)** HEK293T cells expressing circular biSiRA or Broccoli (control) were incubated with bivalent **SiR₂** (500 nM) or monomeric SiR-PEG₂-NH₂ (1.0 µM), originally developed for imaging SiRA aptamers in bacteria[13]. While **SiR₂** enabled imaging biSiRA with excellent signal-to-background ratios in live cells, SiR-PEG₂-NH₂ caused a significant background fluorescence in control cells. Images were shown using the same contrast settings and transfected cells were found by co-transfected mEGFP. Scale bars, 10 µm. **b)** Supplementing the imaging medium with 5 mM magnesium sulfate increased the fluorescence of biSiRA:**SiR₂** in HEK293T cells. **c)** The fluorescence intensity (mean ± s.d) in cells (*N* ≥ 30 cells) described in panel **b**.

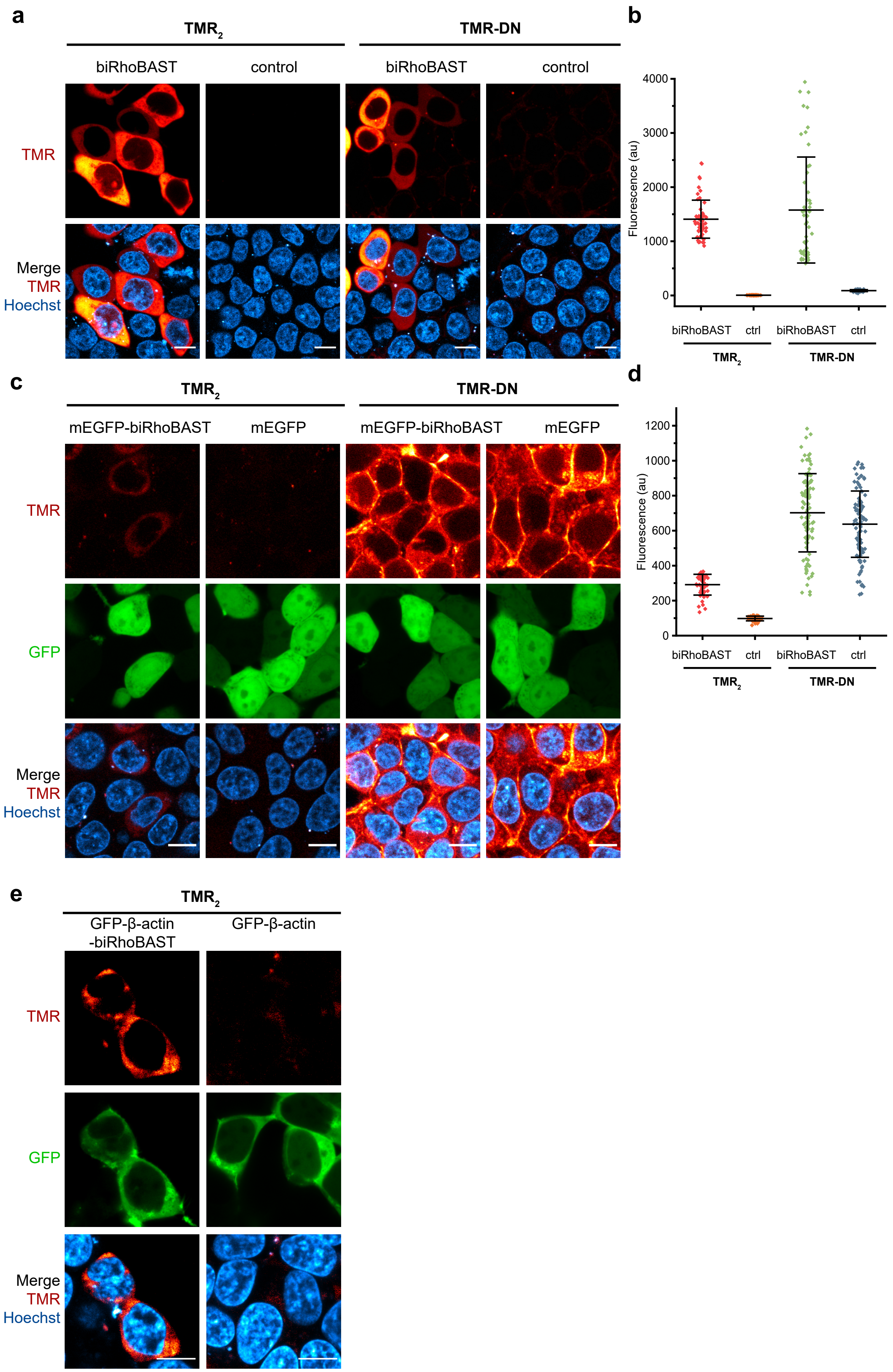
a
TMR2
TMR-DN
biRhoBAST
control
biRhoBAST
control
TMR
Merge
TMR
Hoechst
b
Fluorescence (au)
4000
3000
2000
1000
0
biRhoBAST
ctrl
biRhoBAST
ctrl
TMR2
TMR-DN
c
TMR2
TMR-DN
mEGFP-biRhoBAST
mEGFP
mEGFP-biRhoBAST
mEGFP
TMR
GFP
Merge
TMR
Hoechst
d
Fluorescence (au)
1200
1000
800
600
400
200
0
biRhoBAST
ctrl
biRhoBAST
ctrl
TMR2
TMR-DN
e
TMR2
GFP-β-actin
-biRhoBAST
GFP-β-actin
TMR
GFP
Merge
TMR
Hoechst

**Supplementary Figure 11: $TMR_2$** performs better than TMR-DN for imaging low abundant RNAs in live cells owing to its low background fluorescence.

**a)** HEK293T cells expressing circular biRhoBAST or Broccoli (control) were incubated with either **$TMR_2$** (50 nM) or TMR-DN (100 nM)[14] and imaged. While both probes enabled imaging circular biRhoBAST with remarkable signal-to-background ratios, **$TMR_2$** showed lower background fluorescence in control cells expressing Broccoli aptamer. Scale bars, 10 µm. **b)** The fluorescence intensity (mean ± s.d) in the cytosolic region of cells ($N \geq 50$ cells) described in panel **a**. **c)** HEK293T cells expressing *mEGFP-biRhoBAST* or *mEGFP* mRNA were incubated with either **$TMR_2$** (50 nM) or TMR-DN (100 nM) and imaged. Since the expression level of *mEGFP-biRhoBAST* mRNA is significantly lower than that of circular biRhoBAST, higher laser power was used in this experiment. **$TMR_2$** clearly enabled specific labeling of *mEGFP-biRhoBAST* mRNA in live cells with a signal-to-background ratio of 3. Whereas, TMR-DN did not yield a sufficient signal-to-background ratio to image *mEGFP-biRhoBAST* mRNA in this condition mainly due to the high background fluorescence. Scale bars, 10 µm. **d)** The fluorescence intensity (mean ± s.d) in the cytosolic region of cells ($N \geq 50$ cells) described in panel **c**. **e)** HEKT293T cells expressing *GFP*-β-*actin* mRNA from a chicken-β-actin promoter ($P_{C\beta A}$) with a single biRhoBAST in the 3'UTR were incubated with **$TMR_2$** (50 nM) and imaged with sufficient signal-to-background ratios. Control cells expressing *GFP*-β-*actin* mRNA without biRhoBAST showed minimal background fluorescence after incubation with **$TMR_2$** (50 nM). Scale bar, 10 µm.

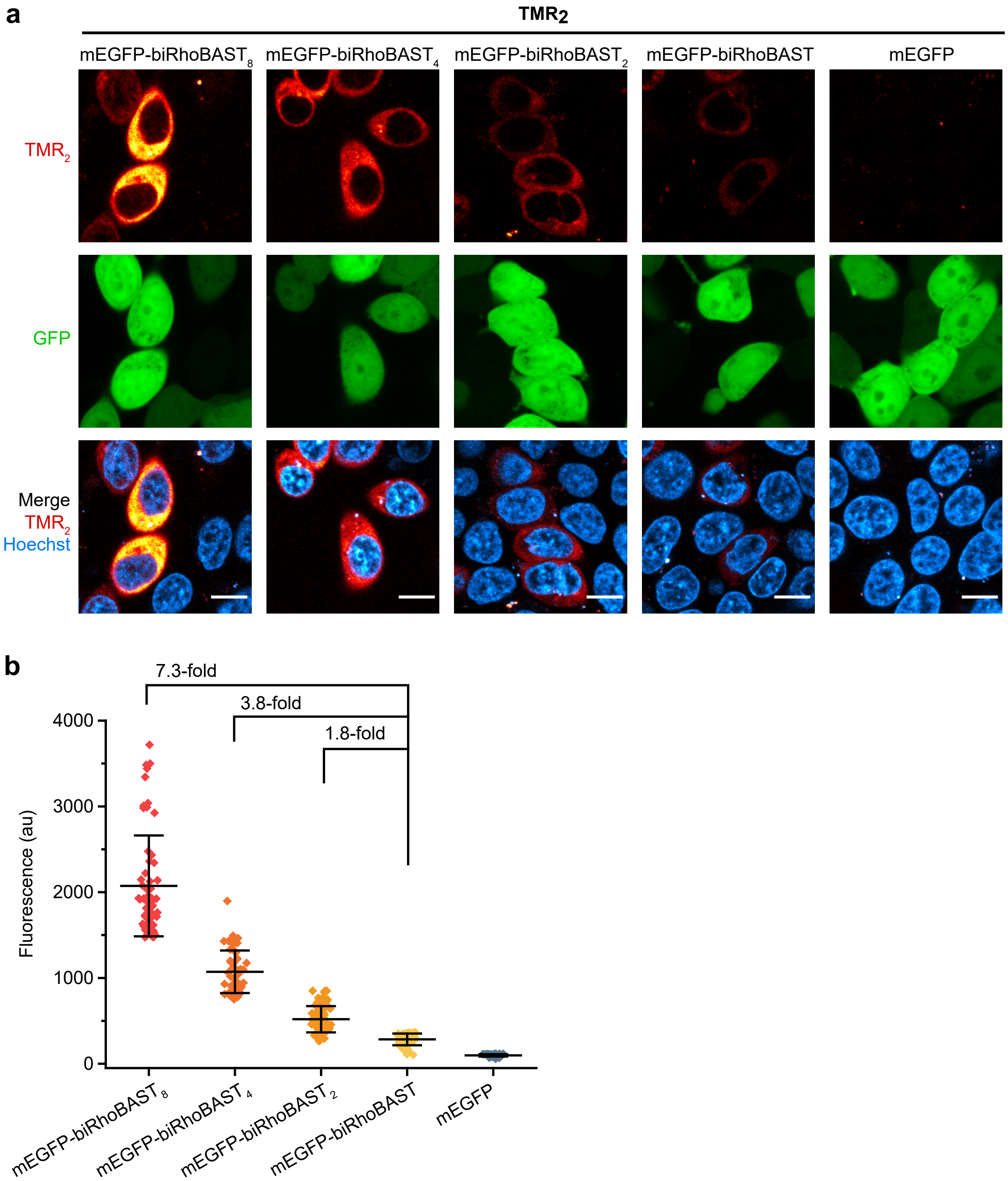


**Supplementary Figure 12:** Multiple copies of biRhoBAST increase the fluorescence signal in live cell RNA imaging.

**a)** Confocal images of HEK293T cells expressing *mEGFP*, *mEGFP-biRhoBAST*, *mEGFP-biRhoBAST*$_2$, *mEGFP-biRhoBAST*$_4$ or *mEGFP-biRhoBAST*$_8$ mRNAs in the presence of **TMR**$_2$ (50 nM). biRhoBAST tags were genetically fused to the 3′ UTR of mEGFP. In the constructs with multiple biRhoBAST units, two synonymous versions of biRhoBAST were used in an alternating order to promote correct folding. Scale bars, 10 µm. **b)** The fluorescence intensity (mean ± s.d) in the cytosolic region of cells ($N \geq 50$ cells) described in panel **a**. For constructs containing 2, 4 and 8 repeats of biRhoBAST, the fluorescence intensity increase relative to mEGFP-biRhoBAST is also shown.

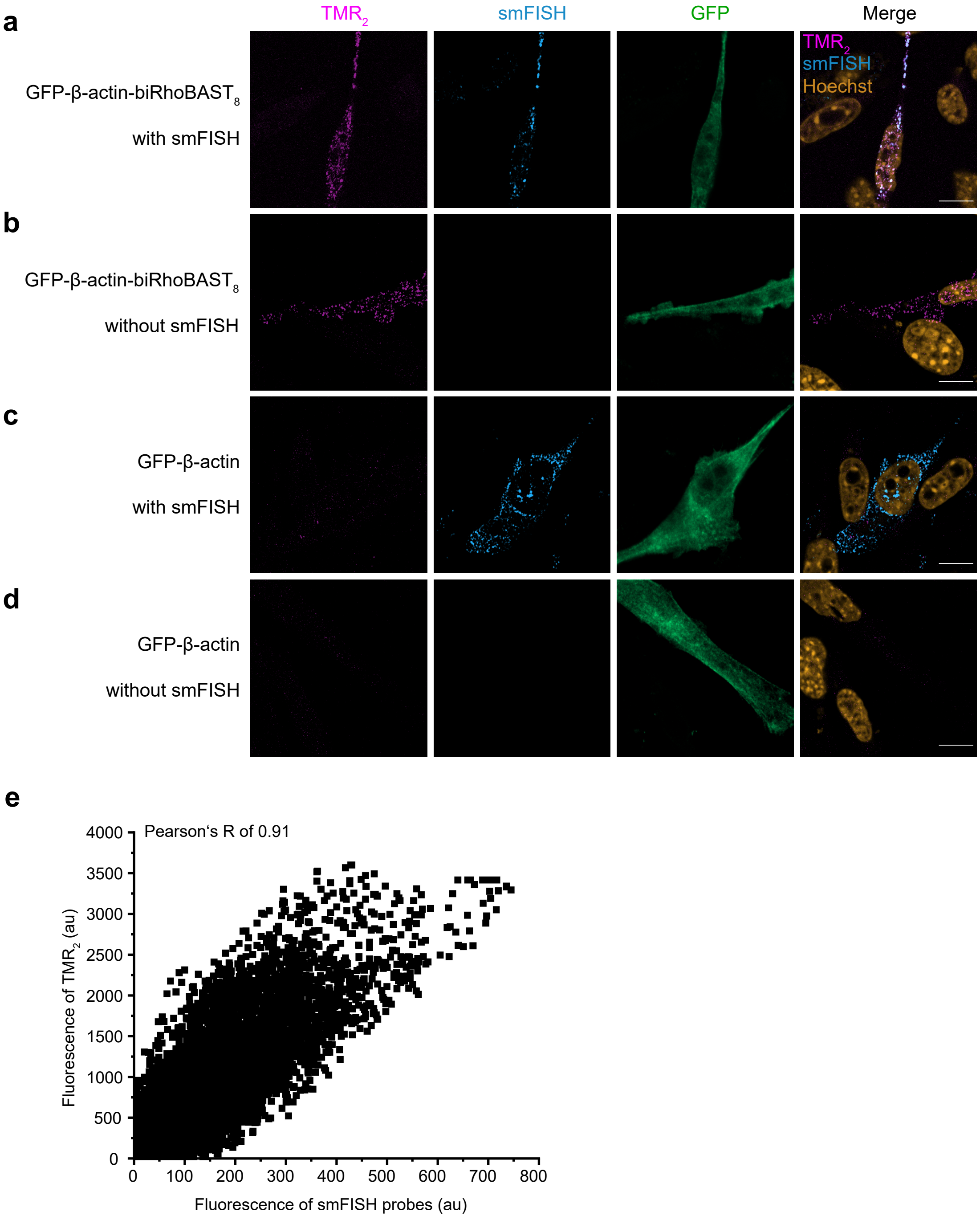


**Supplementary Figure 13:** Validation of the full length labeling and correct localization of *GFP-β-actin-biRhoBAST*$_8$ mRNA by smFISH.

Confocal images of fixed MEF cells expressing **a)** *GFP*-β-*actin-biRhoBAST*$_8$ mRNA after incubation with smFISH probes (targeting the GFP sequence) and **TMR**$_2$ (50 nM); **b)** *GFP*-β-*actin-biRhoBAST*$_8$ mRNA in the presence of **TMR**$_2$ (50 nM); **c)** *GFP*-β-*actin* mRNA after incubation with smFISH probes and **TMR**$_2$ (50 nM); **d)** *GFP*-β-*actin* mRNA in the presence of **TMR**$_2$ (50 nM). Similar localization pattern of mRNAs shown in panels **a** and **c** imply that biRhoBAST$_8$ did not alter the localization of *GFP*-β-*actin* mRNA. Scale bars, 5 µm. **e)** Colocalization test of smFISH probes and **TMR**$_2$ using JACoP[15]. A Pearson's R value of 0.91 was obtained for the positive cell shown in panel **a**, indicating that the majority of the mRNA was intact as a full-length transcript and **TMR**$_2$ signals are not originated from 3' UTR decay fragments.

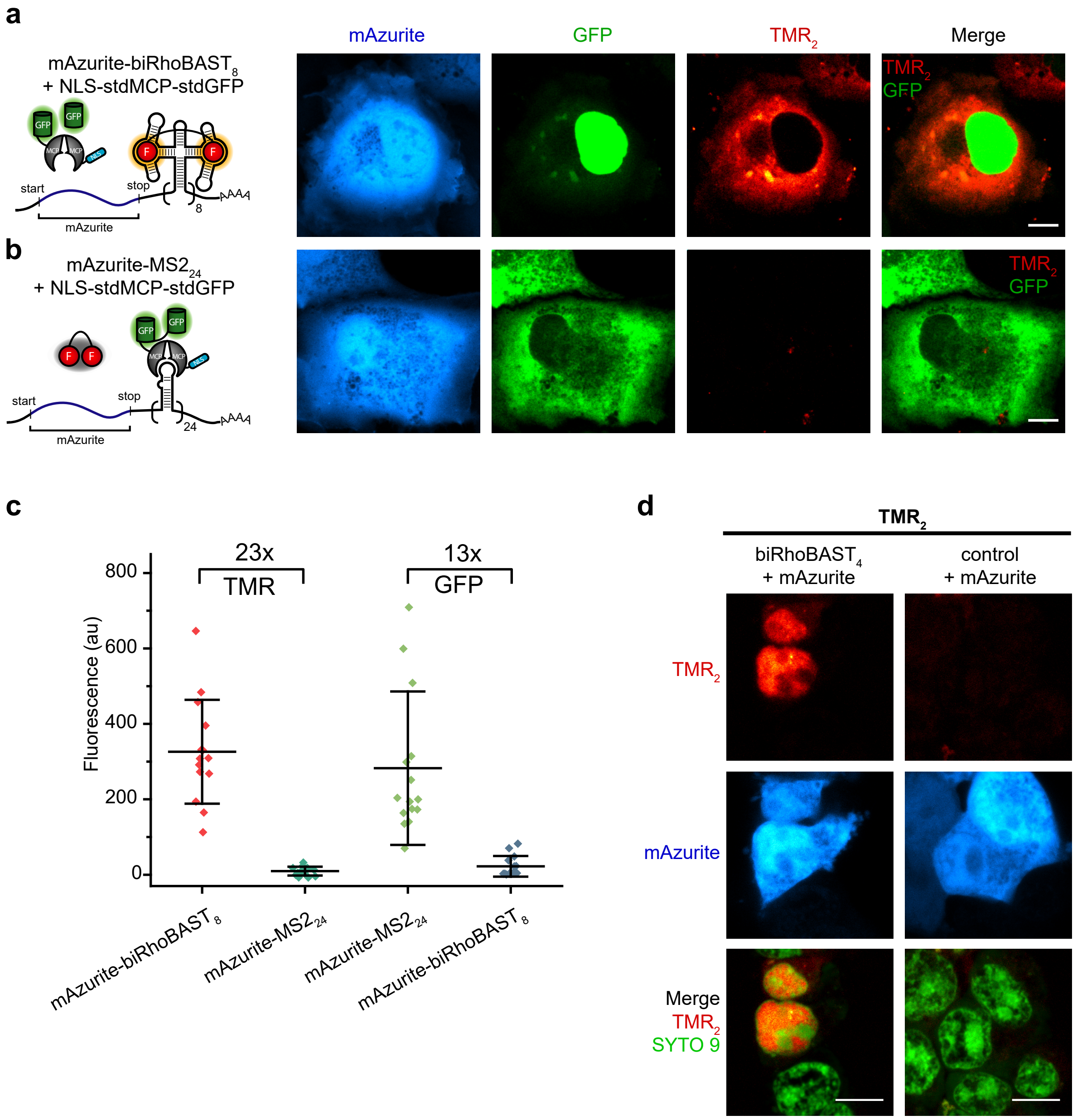


**Supplementary Figure 14**: biRhoBAST:**TMR$_2$** outperforms MS2:MCP-GFP in live cell RNA imaging.

Confocal images of live Cos7 cells expressing **a)** *mAzurite-biRhoBAST$_8$* mRNA and MCP-GFP protein in the presence of **TMR$_2$** (50 nM); **b)** *mAzurite-MS2$_{24}$* mRNA and MCP-GFP protein in the presence of **TMR$_2$** (50 nM). While *mAzurite* mRNA constructs were transcribed under the control of CMV promoter ($P_{CMV}$); MCP-GFP was expressed from the plasmid $P_{UbC}$-NLS-stdMCP-stdGFP with a nuclear localization signal (NLS) at the N-terminus. Due to the NLS sequence, excess MCP-GFP accumulates in the nucleus, resulting in high background fluorescence and making RNA imaging in nucleus very difficult. On the other hand, **TMR$_2$** does not display substantial fluorescence in any particular organelle; therefore, the biRhoBAST:**TMR$_2$** system can be used to image RNAs in different subcellular structures of the cell. Scale bar, 10 µm. **c)** The GFP and TMR fluorescence intensities (mean ± s.d) in cells ($N \geq 13$ cells) described in panel **a** and **b**. biRhoBAST:**TMR$_2$** displayed a higher signal-to-background ratio than MS2:MCP-GFP despite the lower number of chromophores used to label mRNA (16 TMR versus 48 GFP). **d)** biRhoBAST enables imaging nuclear localized RNAs. biRhoBAST was expressed in HEK293T cells under the control of $P_{U6}$ using the plasmid system pAV-U6+27-biRhoBAST$_4$. Cells were also co-transfected with the mAzurite plasmid to easily discriminate biRhoBAST$_4$ expressing cells from un-transfected cells and the images were acquired in the presence of **TMR$_2$** (50 nM). Considering its ability to label nuclear as well as cytosolic RNA, biRhoBAST:**TMR$_2$** system is more versatile than the MS2 system that requires NLS to minimize the cytosolic background. Scale bars, 10 µm.

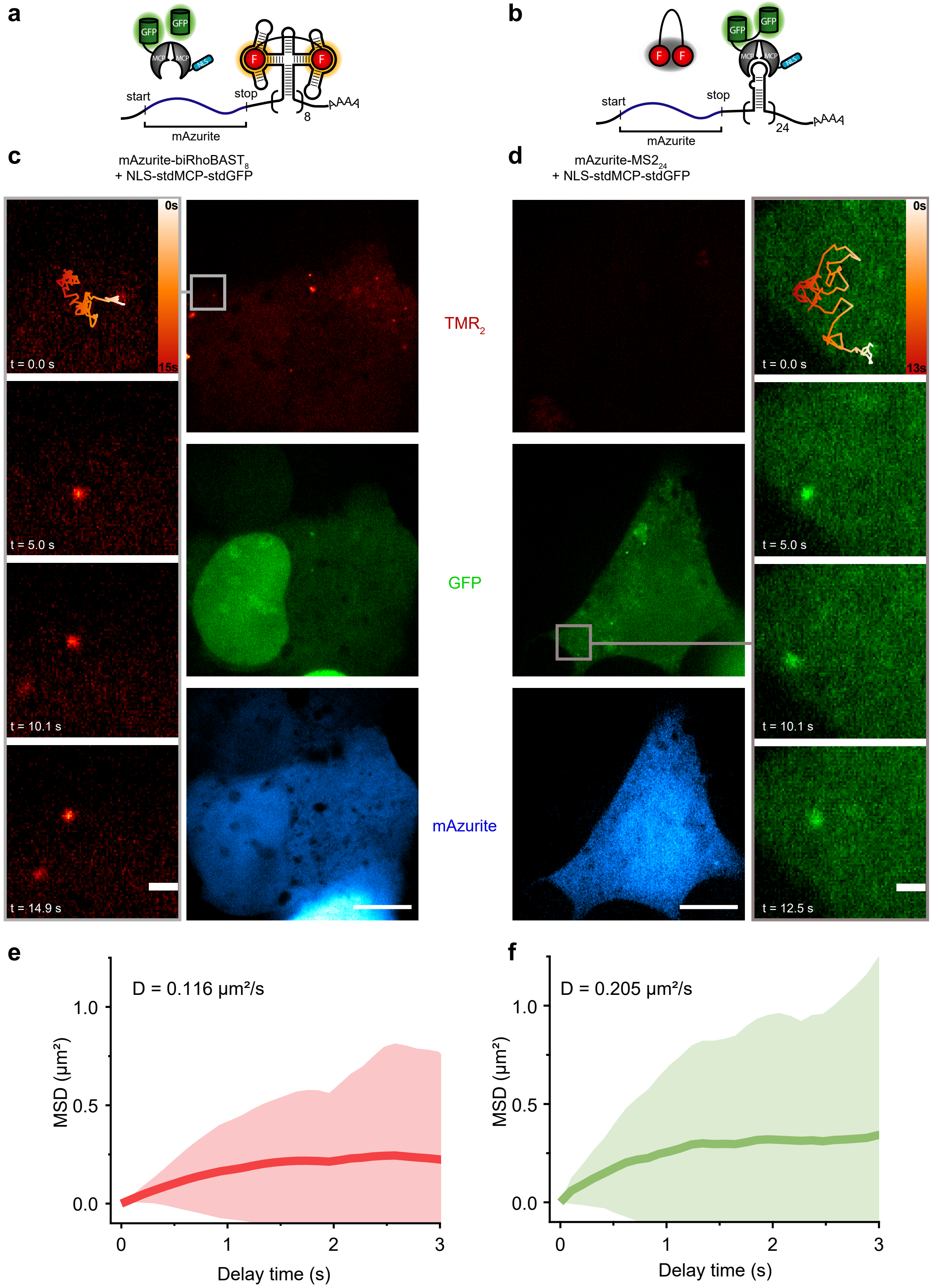


**Supplementary Figure 15:** biRhoBAST:**TMR₂** enables single molecule mRNA tracking in live cells.

Illustration of mRNA constructs **a)** *mAzurite-biRhoBAST₈* and **b)** *mAzurite-MS2₂₄* for single molecule RNA imaging. Tracking *mAzurite* mRNA with 10 frames per second in live U2OS cells with **c)** biRhoBAST₈ and **d)** MS2₂₄ tags. The zoom-ins show an exemplary extracted trajectory at different time points of the acquired data. See also Supplementary Video 2 and 3. Mean square displacements for **e)** *mAzurite-biRhoBAST*$_8$ particles (*N* = 21) and **f)** *mAzurite-MS2*$_{24}$ particles (*N* = 15) were plotted against the delay time for each construct and the diffusion coefficients were calculated by MSDanalyzer[16]. Both biRhoBAST- and MS2-based systems revealed comparable diffusion coefficients for *mAzurite* mRNA, in good agreement with the literature-reported values[17].

**Supplementary Table 1:** Dissociation constants ($K_D$) between the bivalent fluorophore dyes and variants of RhoBAST and SiRA aptamers.

| Dye | Aptamer | $K_D$ (nM) |
|---|---|---|
| | RhoBAST | >46 000 |
| **1** | Linear dimeric RhoBAST | 49 ± 4.0 |
| | biRhoBAST | 0.093 ± 0.007 |
| | RhoBAST | 1813 ± 1276 |
| **2** | Linear dimeric RhoBAST | 6.0 ± 0.5 |
| | biRhoBAST | 0.061 ± 0.010 |
| | RhoBAST | 902 ± 158 |
| **3 ($TMR_2$)** | Linear dimeric RhoBAST | 4.2 ± 0.5 |
| | biRhoBAST | 0.040 ± 0.005 |
| | SiRA | 2975 ± 138 |
| **$SiR_2$** | Linear dimeric SiRA | 15.5 ± 0.8 |
| | biSiRA | 0.291 ± 0.030 |

**Supplementary Table 2:** Photophysical properties of fluorophores and aptamer:fluorophore complexes.

| FLAP | $\lambda_{abs}^{max}$ (nm) | $\lambda_{exc}^{max}$ (nm) | $\lambda_{em}^{max}$ (nm) | QY⁺ | $\varepsilon_{abs}^{max}$ ($M^{-1}cm^{-1}$) | $\varepsilon_{exc}^{max}$* ($M^{-1}cm^{-1}$) |
|---|---|---|---|---|---|---|
| **biRhoBAST:$TMR_2$** | 564 | 564 | 590 | 0.88 | 165 000 | |
| **$TMR_2$** | 517 | 553 | 583 | 0.025 | 136 000 | 51 000 |
| **biRhoBAST:2** | 564 | 564 | 590 | 0.81 | 159 000 | |
| **2** | 517 | 553 | 583 | 0.008 | 115 000 | 56 000 |
| **biRhoBAST:1** | 564 | 564 | 590 | 0.82 | 176 000 | |
| **1** | 517 | 553 | 583 | 0.001 | 160 000 | 48 000 |
| **biSiRA:$SiR_2$** | 649 | 649 | 665 | 0.98 | 200 000 | |
| **$SiR_2$** | 649 | 649 | 670 | 0.15 | 16 000 | |

⁺determined using Pyronin Y (TMR) or Oxazine 1 (SiR) as a reference

*excitation maximum does not match the absorption maximum for the dyes that are mainly in H-dimerized form

**Supplementary Table 3:** Association ($k_{on}$) and dissociation ($k_{off}$) rate coefficients for the complexes between biAptamers and bivalent dyes.

| FLAP | $k_{on}$ ($M^{-1}s^{-1}$) | $k_{off, calc}$ ($s^{-1}$) | $K_D$ (pM) |
|---|---|---|---|
| **biRhoBAST:1** | (6.74 ± 0.18) ×$10^{6}$ | (6.27 ± 0.50) ×$10^{-4}$ | 93 ± 7 |
| **biRhoBAST:2** | (8.99 ± 0.29) ×$10^{6}$ | (5.48 ± 0.92) ×$10^{-4}$ | 61 ± 10 |
| **biRhoBAST:$TMR_2$** | (9.39 ± 0.10) ×$10^{6}$ | (3.76 ± 0.47) ×$10^{-4}$ | 40 ± 5 |
| **biSiRA:$SiR_2$** | (1.73 ± 0.03) ×$10^{5}$ | (5.05 ± 0.53) ×$10^{-5}$ | 291 ± 3 |

**Supplementary Table 4:** Photophysical and biophysical characteristics of our avidity-based systems and other literature reported FLAP systems.

| **FLAP** | **$\lambda_{exc}$ (nm)** | **$\lambda_{em}$ (nm)** | **$\varepsilon$ ($M^{-1}cm^{-1}$)** | **QY** | **Relative brightness*** | **Turn-on (n-fold)** | **$K_D$ (nM)** | **$k_{on}$ ($M^{-1}s^{-1}$)** | **$k_{off}$ ($s^{-1}$)** |
|---|---|---|---|---|---|---|---|---|---|
| **biRhoBAST:$TMR_2$** | 564 | 590 | 164 700 | 0.88 | 2.6 | 55 | 0.040 | $9.39\times10^{6}$ | 0.0004 |
| **biRhoBAST:2** | 564 | 590 | 158 600 | 0.81 | 2.3 | 92 | 0.061 | $8.99\times10^{6}$ | 0.0005 |
| **biRhoBAST:1** | 564 | 590 | 175 900 | 0.82 | 2.6 | 284 | 0.093 | $6.74\times10^{6}$ | 0.0006 |
| **biSiRA:$SiR_2$** | 649 | 663 | 200 000 | 0.98 | 3.6 | 100 | 0.291 | $1.73\times10^{5}$ | $5.05\times10^{-5}$ |
| **RhoBAST:TMR-DN[14]** | 564 | 590 | 96 000 | 0.57 | 1.0 | 26 | 15 | $1.6\times10^{8}$ | 2.94 |
| **SiRA:SiR-$PEG_2$-$NH_2$[13]** | 649 | 662 | 86 000 | 0.98 | 1.5 | 7 | 430 | $6.6\times10^{3}$ | 0.0028 |
| **Riboglow:Cbl-Atto590[12]** | 594 | 622 | 120 000 | 0.31 | 0.7 | 5 | 34 | - | - |
| **Riboglow:Cbl-Cy5[12]** | 646 | 666 | 271 000 | 0.25 | 1.2 | 2.7 | 3 | - | - |
| **Broccoli:BI[10]** | 470 | 505 | 33 600 | 0.67 | 0.4 | 1 100 | 51 | $1.4\times10^{4}$ | 0.0006 |
| **MangoII:TO1-Biotin[18]** | 510 | 535 | 77 500 | 0.21 | 0.3 | 1 300 | 0.7 | - | - |
| **DIR2:DIR-Pro[19]** | 600 | 658 | 164 000 | 0.33 | 1.0 | 80 | 252 | - | - |
| **o-Coral:Gemini-561[2]** | 580 | 596 | 141 000 | 0.58 | 1.5 | 13 | 73 | - | - |
| **Pepper:HBC620[9]** | 577 | 620 | 100 000 | 0.58 | 1.1 | 12 600 | 6.1 | $5.5\times10^{4}$ | 0.0041 |

* calculated as QY × $\varepsilon_{exc}^{max}$ and normalized to the brightness of RhoBAST:TMR-DN.

**Supplementary Table 5:** DNA sequences used in this study.

| | |
|---|---|
| T7 promoter (ϕ 6.5) (T7P) | TAATACGACTCACTATA |
| RhoBAST | T7P- GGAACCTCCGCGAAAGCGGTGAAGGAGAGGCGCAAGGTTAACCGCCTCAGGTTCC |
| biRhoBAST linear | T7P- GGAACCTCCGCGAAAGCGGTGAAGGAGAGGCGCAAGGTTAACCGCCTCAGGTTCC AAGAAAAGAAAAGGCCTCCGCGAAAGCGGTGAAGGAGCGCCACAAGGTTAACTGGCGCAGGCCTT |
| biRhoBAST | T7P- GGTATGTGGGAGGAACCTCCGCGAAAGCGGTGAAGGAGAGGCGCAAGGTTAACCGCCTCAGGTTCCAGGAGGATTCGTCCTCCAAAGGCCTCCGCGAAAGCGGTGAAGGAGCGCCACAAGGTTAACTGGCGCAGGCCTTACCCACATACC |
| biRhoBAST (V1) | taatctagaGGTATGTGGGAGGAACCTCCGCGAAAGCGGTGAAGGAGAGGCGCAAGGTTAACCGCCTCAGGTTCCAGGAGGATTCGTCCTCCAAAGGCCTCCGCGAAAGCGGTGAAGGAGCGCCACAAGGTTAACTGGCGCAGGCCTTACCCACATACCgggccctat |
| pcDNA3 Nhe1 fwd | gctagctattctatagtgtcacctaaatgctagagctcgc |
| pcDNA3 Nhe1 rev | ggtatgtgggtaaggcctgcgc |
| biRhoBAST (V2) | atctctagaGGAGCCCGGAAGGAACCTCCGCGAAAGCGGTGAAGGAGAGGCGCAAGGTTAACCGCCTCAGGTTCCAGCGGAGTTCGCTCCGCAAAGGCCTCCGCGAAAGCGGTGAAGGAGCGCCACAAGGTTAACTGGCGCAGGCCTTATCCGGGCTCCgctagcgtc |
| mAzurite seq fwd | cgagaagcgcgatcacatgg |
| mAzurite seq rev | ggaggtgtgggaggttttttaa |
| circular-biRhoBAST | agtgcggccgcTGTGGGAGGAACCTCCGCGAAAGCGGTGAAGGAGAGGCGCAAGGTTAACCGCCTCAGGTTCCAGGAGGATTCGTCCTCCAAAGGCCTCCGCGAAAGCGGTGAAGGAGCGCCACAAGGTTAACTGGCGCAGGCCTTACCCACAgtggccgcggtcg |
| mAzu stop fwd | tcgaggggtctagaggttaccgatatccatatggcggccgcatcgata |
| mAzu stop rev | agcttatcgatgcggccgccatatggatatcggtaacctctagacccC |
| MS2v6 seq fwd | ctaggacctgcaggcatgcaag |
| smFISH porbes (HuluFISH) for GFP,<br>19 matching for mAzurite | TCCAGCTCGACCAGGATG, GCTGAACTTGTGGCCGTTTAC, TTCAGGGTCAGCTTGCCGT, CCGGTGGTGCAGATGAAC, TAGGTCAGGGTGGTCACGAGG, CTTGAAGAAGTCGTGCTGC, TTCATGTGGTCGGGGTAGCGG, TCCTGGACGTAGCCTTCG, TGCCGTCGTCCTTGAAG, TCGAACTTCACCTCGGCG, GATGCGGTTCACCAGGGT, AGTCGATGCCCTTCAGCTC, GATGTTGCCGTCCTCCTTGA, AGTTGTACTCCAGCTTGTG, ATAGACGTTGTGGCTGTTG, GTTCTTCTGCTTGTCGGCC, TGTGGCGGATCTTGAAGTTC, ATGCCGAGAGTGATCCCG, TTCTGCTGGTAGTGGTCGGC, TAGTGGTTGTCGGGCAGCAG, CGTTGGGGTCTTTGCTCAG, ACCATGTGATCGCGCTTCT, GTCACGAACTCCAGCAGG |
| SiRA | T7P- GGCCACCGGGTTTGAAAACCTGGCTGCTTCGGCAGTTGTATCCTTTGGCC |
| biSiRA linear | T7P- GGCCACCGGGTTTGAAAACCTGGCTGCTTCGGCAGTTGTATCCTTTGGCCAAGAAAAGAACCCCACCGGGTTTGAAAACCTGGCTGCTTCGGCAGTTGTATCCTTTGGGG |
| biSiRA | T7P- GGTATGTGGGAGGCCACCGGGTTTGAAAACCTGGCTGCTTCGGCAGTTGTATCCTTTGGCCAGGAGGAGAAATCCTCCACCCCACCGGGTTTGAAAACCTGGCTGCTTCGGCAGTTGTATCCTTTGGGGACCCACATACC |
| biSiRA (V1) | ctcagatctcgacgggtctagactcgag GGTATGTGGGAGGCCACCGGGTTTGAAAACCTGGCTGCTTCGGCAGTTGTATCCTTTGGCCAGGAGGAGAAATCCTCCACCCCACCGGGTTTGAAAACCTGGCTGCTTCGGCAGTTGTATCCTTTGGGGACCCACATACC |

| | |
|---|---|
| | gtcgacggt |
| circular-biSiRA | agtgcggccgcTGTGGGAGGCCACCGGGTTTGAAAACCTGGCTGCTTCGGCAGTTGTATCCTTTGGCCAGGAGGAGAAATCCTCCACCCCACCGGGTTTGAAAACCTGGCTGCTTCGGCAGTTGTATCCTTTGGGGACCCACAgtggccgcggtcg |
| circular-Mango2 | agtgcggccgcGCGTACGAAGGAGAGGAGAGGAAGAGGAGAGTACGCgtggccgcggtcg |
| circular-Pepper | agtgcggccgcCCCAATCGTGGCGTGTCGGCCTGCTTCGGCAGGCACTGGCGCCGGgtggccgcggtcg |

**Supplementary Video 1**: Verification of single mRNA tagging with a biRhoBAST-MS2 tandem construct.

Mobile puncta were imaged in Cos7 cells co-expressing MCP-GFP and (i) *mAzurite-biRhoBAST$_8$-MS2$_{24}$* mRNA (ii) *mAzurite-biRhoBAST$_8$* mRNA (iii) *mAzurite-MS2$_{24}$* mRNA were imaged in the presence of **TMR$_2$** (500nM). Exposure time per frame: 100 ms. Scale bar, 5 µm.

**Supplementary Video 2**: Single mRNA tracking with biRhoBAST:**TMR$_2$**.

Single RNA molecules were visualized and tracked in U2OS cells expressing *mAzurite-biRhoBAST$_8$* mRNA in the presence of **TMR$_2$** (500nM). Exposure time per frame: 100 ms. Scale bar, 5 µm.

**Supplementary Video 3**: Single mRNA tracking with MS2:MCP-GFP.

Single RNA molecules were visualized and tracked in U2OS cells co-expressing *mAzurite-MS2$_{24}$* mRNA and MCP-GFP. Exposure time per frame: 100 ms. Scale bar, 5 µm.

#### **Supplementary Note 1:** Quenching mechanism of **SiR₂**.

3-carboxy silicon rhodamines (SiRs) exist in an equilibrium between a colorless spirolacton form and a fluorescent zwitterionic open form. This equilibrium favors either open or closed form depending on the environment and accounts for the observed fluorogenicity of SiR dyes[13, 20]. However, in **SiR$_2$** we believe that the origin of the high fluorogenicity compared to the original SiRA system (100-fold vs 7-fold fluorescent enhancement) is a combination of spirocyclization and H-dimer formation.

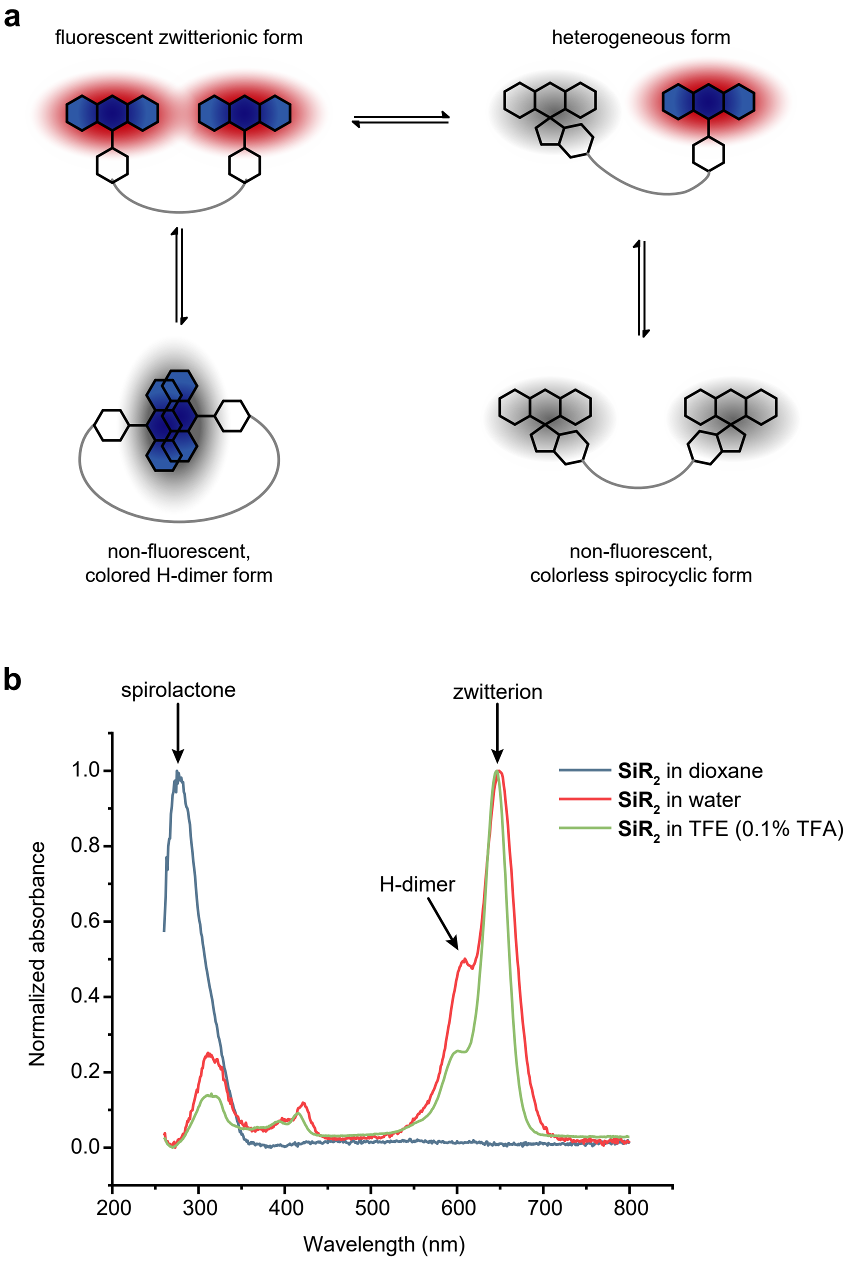


**Supplementary Note Figure I: a)** Scheme displaying various fluorescent and non-fluorescent species of **SiR$_2$** in dynamic equilibrium. Both spirocyclization and H-dimer formation play a role in the quenching mechanism of **SiR$_2$**. **b)** Absorption spectra of **SiR$_2$** (5 µM) in dioxane, water and TFE (with 0.1% TFA) to investigate the spirolactone and H-dimer contribution to the dye, normalized to the maximum absorbance values. In dioxane SiRs are typically in the colorless and non-fluorescent spirolactone form, which we found in the spectrum of **SiR$_2$** in dioxane at 276 nm[20]. TFE supplemented with TFA prevents not only spirolactonization due to acidic pH, but also H-dimer formation[3, 20] due to favourable interactions between the organic solvent and the SiR. Therefore, **SiR$_2$** exists in the highly colored form displaying absorption maximum at 649 nm in TFE (+0.1% TFA). **SiR$_2$** in water also showed the colored zwitterionic form (647 nm), as well as an increased local maximum at 607 nm indicating H-dimers[21] and a band around 311 nm indicating the spirolactone form[20].

The first evidence for the presence of H-dimers can be seen from the absorption spectrum of **$SiR_2$**. Although the absorption spectrum of **$SiR_2$** did not result in a single blue-shifted absorption maximum relative to monomeric SiRs (as observed in **$TMR_2$**), the ratio of the 607 nm band (originated from the H-dimer) to 649 nm band (originated from the open form of SiR) increased by 1.7-fold compared to monomeric SiR-$PEG_2$-$NH_2$. These data suggest the co-presence of H-dimers and open form of SiR, which is in dynamic equilibrium with the closed form

The second indication for the H dimer formation is the relatively low quantum yield of **$SiR_2$** (QY = 0.15) compared to SiR-$PEG_2$-$NH_2$ (QY = 0.51)[13]. We claim that, rather than spirolactonization, an additional quenching effect such as H-dimer formation in **$SiR_2$** can be responsible for this decrease. Since H-dimers are colored but not fluorescent, their presence decreases the fluorescence quantum yield of probes. Moreover, the differences in the QYs of **$SiR_2$** and SiR-$PEG_2$-$NH_2$ cannot be solely explained by the probes' different extent of spirolactonization, as the core fluorophore structure of the closed probes is essentially the same. Also, the presence of closed form does not significantly change the QY of SiR probes, as the closed form is colorless and non-fluorescent.

Finally, we compared the absorption spectra of **$SiR_2$** in different solvents to verify the contribution of H-dimer formation and spirolactonization (Supplementary Note Figure I). In water, **$SiR_2$** shows typical bands that correspond to H-dimers (607 nm) and spirolactones (311 nm). Therefore, the high fluorogenicity of **$SiR_2$** is likely a result of spirolactonization as well as H-dimer formation.

#### **Supplementary Note 2:** Synthesis Route of Fluorophore Dimers

The synthesis route for polyethylene glycol linkers and fluorophore dimers is described in the scheme below. First, commercially available azide **4** was converted into diamine **5** by reduction in methanol catalyzed by palladium on activated charcoal adapting a protocol from Chiba *et al*[22]. Diamine **5** was added to carboxylic acid **6**, which was activated with TSTU for 1 h before mixing, adapting a procedure of Robers *et al*[23]. Boc-NH-groups were then deprotected in TFA/DCM, adapting a protocol from Chiba *et al*[22]. The TMR dimers were synthesized by coupling with TSTU and subsequent addition of the respective diamine linker, adapting a protocol of Robers *et al*[23]. **SiR$_2$** was synthesized by coupling with PyBOP and followed by the addition of diamine **8**, as described by Wirth *et al*[13].

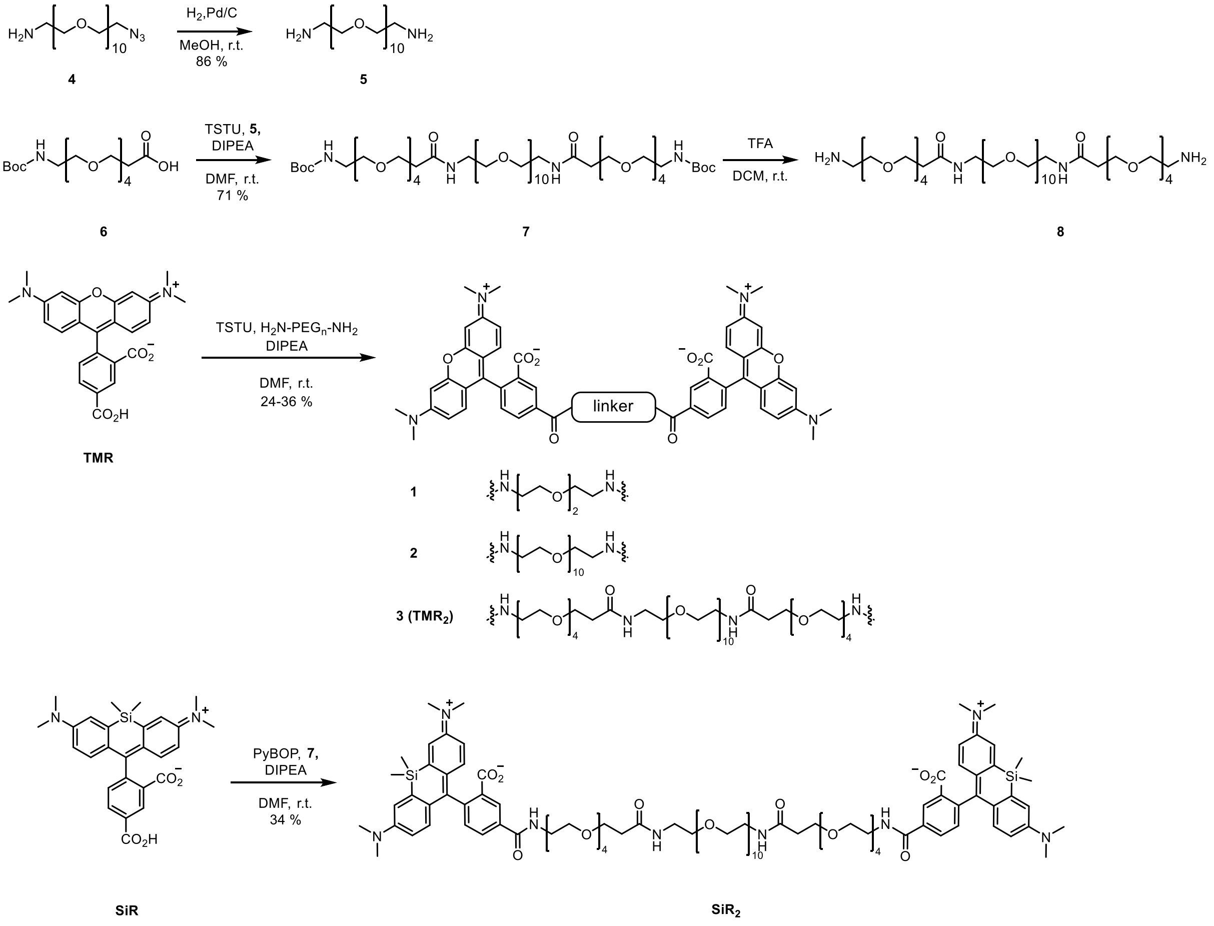

$H_2N$-$PEG_{10}$-$NH_2$ (**5**):

Under argon atmosphere, azide **4** (160.0 mg, 303.8 µmol) was dissolved in 10 mL MeOH and suspended with a spatula tip of 10% palladium on activated charcoal. The argon atmosphere was exchanged to hydrogen atmosphere, the mixture was stirred for 18 h at room temperature and the conversion monitored by TLC. The product was filtered with a 0.46 µm spin filter for 2 min at 5000 rpm. The solvents were removed under reduced pressure to afford the product with a yield of 86% (131.0 mg, 261.7 µmol).

**$R_f$** (25:5:1 v/v $CHCl_3$:MeOH:$NH_4OH_{aq}$(25%)): 0.60.
**HRMS (ESI):** *m/z* $[M+H]^+$ meas. 501.3389, calc. 501.3382.
**$^1$H-NMR (500 MHz, $CD_3OD$)**: $\delta$ = 3.64 (m, 36H), 3.53 (t, *J* = 5.3 Hz, 4H), 2.79 (t, *J* = 5.3 Hz, 4H) ppm.
**$^{13}$C{$^1$H}-NMR (126 MHz, $CD_3OD$)**: $\delta$ = 73.3, 71.5, 41.1 ppm.

Boc-NH-$PEG_{20}$-NH-Boc (**7**)

A solution of **6** (36.5 mg, 99.9 µmol, 2.5 eq), TSTU (32.5 mg, 107.9 µmol, 2.7 eq.) and DIPEA (13.9 mg, 18.8 µL, 107.9 µmol, 2.7 eq.) in 10 mL dry DMF was stirred for 1 h at room temperature. Then, PEG10-diamine (20 mg, 40.0 µmol, 1.0 eq.) was added (dissolved in 0.5 mL dry DMF) and the mixture was stirred for 16 h at room temperature. After evaporating the solvent, the obtained crude product was purified by flash column chromatography on silica gel (Gradient 10:1:0 to 35:5:1 v/v $CHCl_3$:MeOH:$NH_4OH_{aq}$(25%)). The product was obtained with a yield of 71% (34 mg, 28.5 µmol).

**$R_f$** (35:5:1 v/v $CHCl_3$:MeOH:$NH_4OH_{aq}$(25%)): 0.47.
**HRMS (ESI):** *m/z* $[M+H]^+$ meas. 1195.7328, calc. 1195.7270.
**$^1$H-NMR (500 MHz, $CD_3OD$)**: $\delta$ = 3.72 (t, *J* = 6.2 Hz, 4H), 3.68 – 3.57 (m, 60H), 3.54 (t, *J* = 5.6 Hz, 4H), 3.50 (t, *J* = 5.6 Hz, 4H), 3.37 (t, *J* = 5.6 Hz, 4H), 3.21 (t, *J* = 5.7 Hz, 4H), 2.46 (t, *J* = 6.2 Hz, 4H), 1.43 (s, 18H) ppm.
**$^{13}$C{$^1$H}-NMR (126 MHz, $CD_3OD$)**: $\delta$ = 174.0, 71.5, 71.4, 71.3, 71.0, 70.6, 68.3, 41.3, 40.4, 37.6, 28.8 ppm.

$H_2N$-$PEG_{20}$-$NH_2$ (**8**)

To a solution of **7** (34 mg, 28.5 mol) in 3 mL $CH_2Cl_2$, 1 mL TFA was added dropwise at 0 °C. After complete addition, it was stirred at room temperature for 8 h. The solvents were removed under reduced pressure to afford the product which was used without further purification and characterization.

**$R_f$** (35:5:1 v/v $CHCl_3$:MeOH:$NH_4OH_{aq}$(25%)):0.26.
**HRMS (ESI):** *m/z* $[M+H]^+$ meas. 995.6239, calc. 995.6221.

General procedure for TMR-based Dimers

TMR (3.0 eq.) was activated with TSTU (3.2 eq.) and DIPEA (3.2 eq.) in dry DMF for 1 h at room temperature. Then, $H_2N$-$PEG_n$-$NH_2$ (1.0 eq.) was added and the reaction stirred for 4-16 h at room temperature until TLC reaction control showed no further conversion. The solvents were removed under reduced pressure and the product purified by flash column chromatography on silica gel (25:5:1

v/v $CHCl_3$:MeOH:$NH_4OH_{aq}$(25%)) followed by reverse phase HPLC purification. Yields can be found in the following.

TMR-PEG$_2$-TMR (**1**):

With 1,8-diamino-3,6-dioxaoctane and 80.0 mg (185.9 µmol) TMR, the product was obtained with a yield of 34% (20.6 mg, 21.2 µmol), following general procedure for TMR based dimers.

**R$_f$** (25:5:1 v/v $CHCl_3$:MeOH:$NH_4OH_{aq}$(25%)): 0.56.
**HPLC**: Gradient (25% to 45% ACN in water with 0.1% TFA in 40 min) Elution time 34.4 min.
**HRMS (ESI):** *m/z* $[M+H]^+$ meas. 973.4140, calc. 973.4131.
**$^1$H-NMR (500 MHz, $CD_3OD$)**: $\delta$ = 8.76 (d, $J$ = 1.8 Hz, 2H), 8.23 (dd, $J$ = 7.9, 1.9 Hz, 2H), 7.47 (d, $J$ = 7.9 Hz, 2H), 7.10 (d, $J$ = 9.4 Hz, 4H), 7.00 (dd, $J$ = 9.4, 2.4 Hz, 4H), 6.97 (d, $J$ = 2.4 Hz, 4H), 3.78 (t, $J$ = 5.4 Hz, 4H), 3.76 (s, 4H), 3.68 (t, $J$ = 5.4 Hz, 4H), 3.32 (s, 24H) ppm.
**$^{13}$C{$^1$H}-NMR (126 MHz, $CD_3OD$)**: $\delta$ = 168.2, 167.6, 160.6, 159.0, 158.9, 138.0, 137.6, 132.2, 132.0, 131.8, 131.3, 115.5, 114.7, 97.4, 71.5, 70.6, 41.2, 40.9 ppm.

TMR-PEG$_{10}$-TMR (**2**):

With PEG10-diamine **5** and 80.0 mg (185.9 µmol) TMR ,the product was obtained with a yield of 36% (29.6 mg, 22.3 µmol), following general procedure for TMR based dimers.

**R$_f$** (25:5:1 v/v $CHCl_3$:MeOH:$NH_4OH_{aq}$(25%)): 0.45.
**HPLC:** Gradient (30% to 50% ACN in water with 0.1% TFA in 40 min) Elution time 21.3 min.
**HRMS (ESI):** *m/z* $[M+H]^+$ meas. 1325.6183, calc. 1325.6228.
**$^1$H-NMR (500 MHz, $CD_3OD$)**: $\delta$ = 8.78 (d, $J$ = 1.8 Hz, 2H), 8.28 (dd, $J$ = 7.9, 1.8 Hz, 2H), 7.53 (d, $J$ = 7.9 Hz, 2H), 7.13 (d, $J$ = 9.5 Hz, 4H), 7.05 (dd, $J$ = 9.5, 2.5 Hz, 4H), 6.98 (d, $J$ = 2.5 Hz, 4H), 3.57-3.74 (m, 44H), 3.31 (s, 24H) ppm.
**$^{13}$C{$^1$H}-NMR (126 MHz, $CD_3OD$)**: $\delta$ = 168.2, 167.4, 160.6, 159.0, 158.9, 138.1, 137.7, 132.9, 132.4, 131.9, 131.4, 115.6, 114.7, 97.5, 71.6, 71.5, 71.4, 70.5, 41.3, 40.9 ppm.

TMR-PEG$_{20}$-TMR (**3** or **TMR$_2$**):

With PEG20-diamine **8** and 18.4 mg (42.8 µmol) TMR, the product was obtained with a yield of 24% (3.0 mg, 1.65 µmol), following general procedure for TMR based dimers.

**R$_f$** (25:5:1 v/v $CHCl_3$:MeOH:$NH_4OH_{aq}$(25%)): 0.63.
**HPLC**: 36% ACN in water with 0.1% TFA. Elution time 24.6 min.
**HRMS (ESI):** *m/z* $[M+Na]^+$ meas. 1842.8933, calc. 1842.8920.
**$^1$H-NMR (500 MHz, $CD_3OD$)**: $\delta$ = 8.76 (d, $J$ = 1.9 Hz, 2H), 8.26 (dd, $J$ = 7.9, 1.8 Hz, 2H), 7.52 (d, $J$ = 7.8 Hz, 2H), 7.16 (d, $J$ = 9.5 Hz, 4H), 7.06 (dd, $J$ = 9.5, 2.5 Hz, 4H), 6.98 (d, $J$ = 2.4 Hz, 4H), 3.97 – 3.42 (m, 80H), 2.42 (t, $J$ = 6.2 Hz, 4H) ppm.
**$^{13}$C{$^1$H}-NMR (126 MHz, $CD_3OD$)**: $\delta$ = 173.9, 168.3, 160.7 159.1, 159.0, 138.0, 137.7, 132.1, 132.0, 131.8, 131.2, 115.5, 114.7, 97.5, 71.6, 70.6, 68.3, 41.3, 41.0, 37.6 ppm.

SiR-PEG20-SiR (**$SiR_2$**)

Under argon atmosphere, 35 mg PyBOP (67.3 µmol, 2.80 eq.), 15 µL DIPEA (11.1 mg, 86.0 µmol, 3.60 eq.) and 25 mg 5C-SiR (52.9 µmol, 2.20 eq.) were dissolved in 2 mL dry DMF and stirred for 1 h at room temperature. Then, **8** (23.9 mg ,24.0 µmol, 1.00 eq.) in 0.5 mL dry DMF was added dropwise and the reaction mixture stirred at room temperature overnight. The solvents were removed under reduced pressure and the product purified by column chromatography (gradient 3% MeOH in DCM to 10% MeOH), followed by semi-preparative reverse phase HPLC. The product was obtained as a blue solid with a yield of 34% (15.6 mg, 8.17 µmol).

**$R_f$** (5% MeOH in DCM): 0.21.
**HPLC**: Gradient (30% to 60% ACN in water with 0.1% TFA in 40 min). Elution time 24 min.
**HRMS (ESI):** *m/z* $[M+H]^+$ meas. 1903.9677, calc. 1903.9647.
**$^1$H-NMR (500 MHz, $CD_3OD$):** δ = 8.53 (s, 1H), 8.20 (d, *J* = 8.0 Hz, 1H), 7.38 (d, *J* = 8.0, 1H), 7.16 (d, *J* = 2.8 Hz, 2H), 6.81 (d, *J* = 9.2 Hz, 2H), 6.67 (dd, *J* = 9.2, 2.9 Hz, 2H), 3.72-3.49 (m, 80H), 3.10 (s, 24H), 2.41 (t, *J* = 6.2 Hz, 2H), 0.64 (s, 6H), 0.56 (s, 6H) ppm.

**NMR spectra of products**

$^1$H NMR (500 MHz, $CD_3OD$)

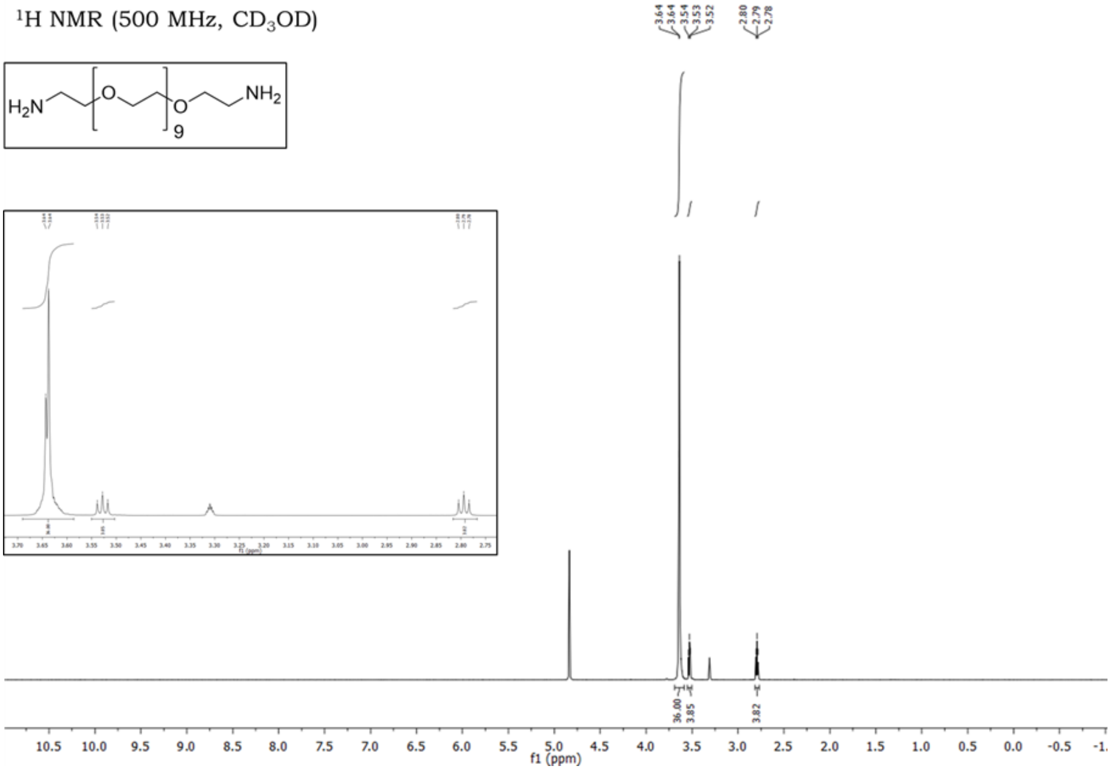


$^{13}$C NMR (126 MHz, $CD_3OD$)

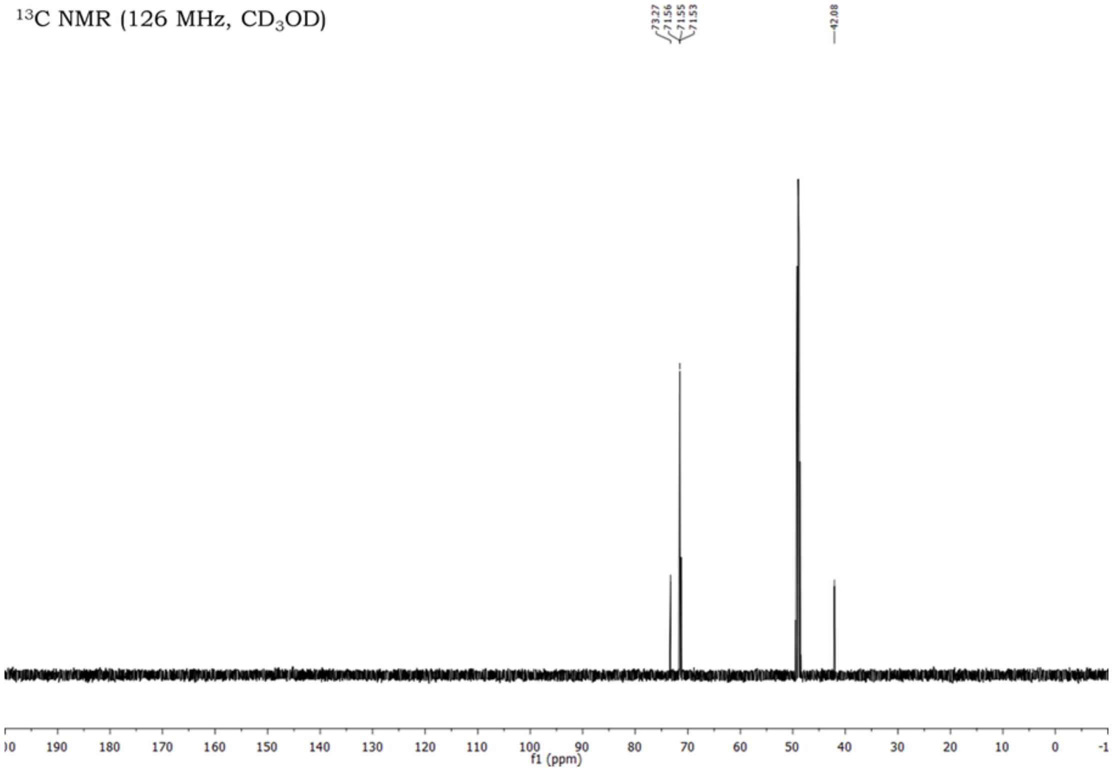

$^{1}H$ NMR (500 MHz, $CD_3OD$)

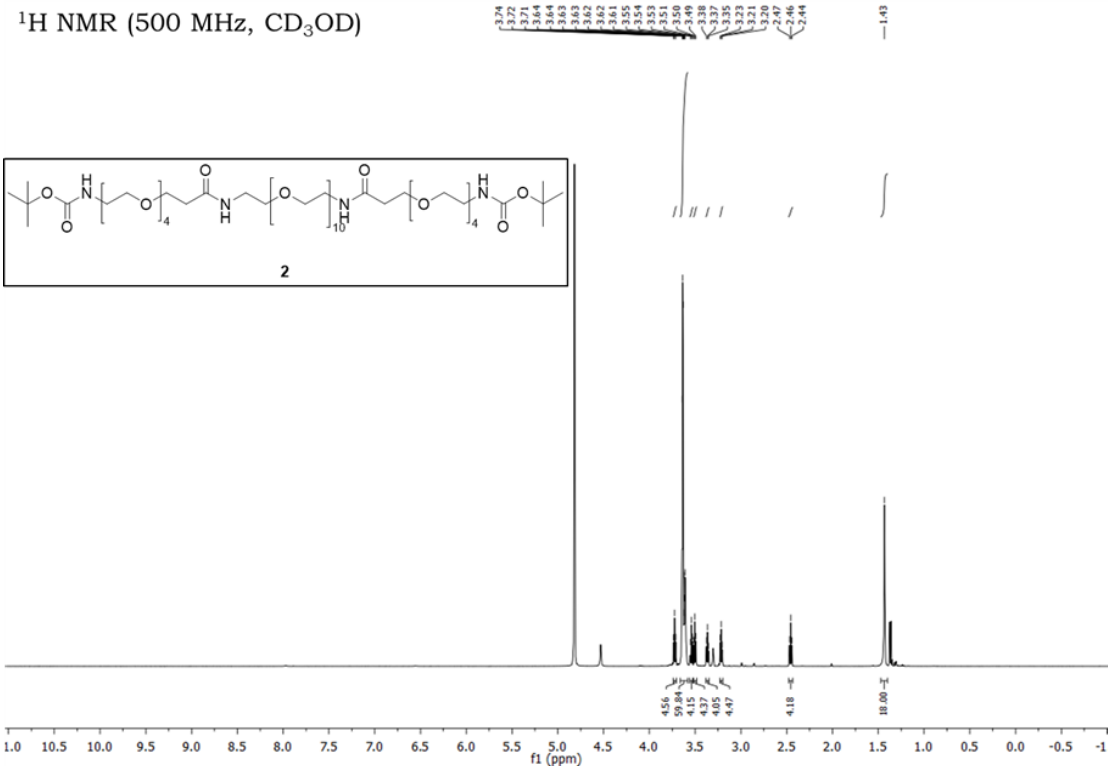

2

$^{13}C$ NMR (126 MHz, $CD_3OD$)

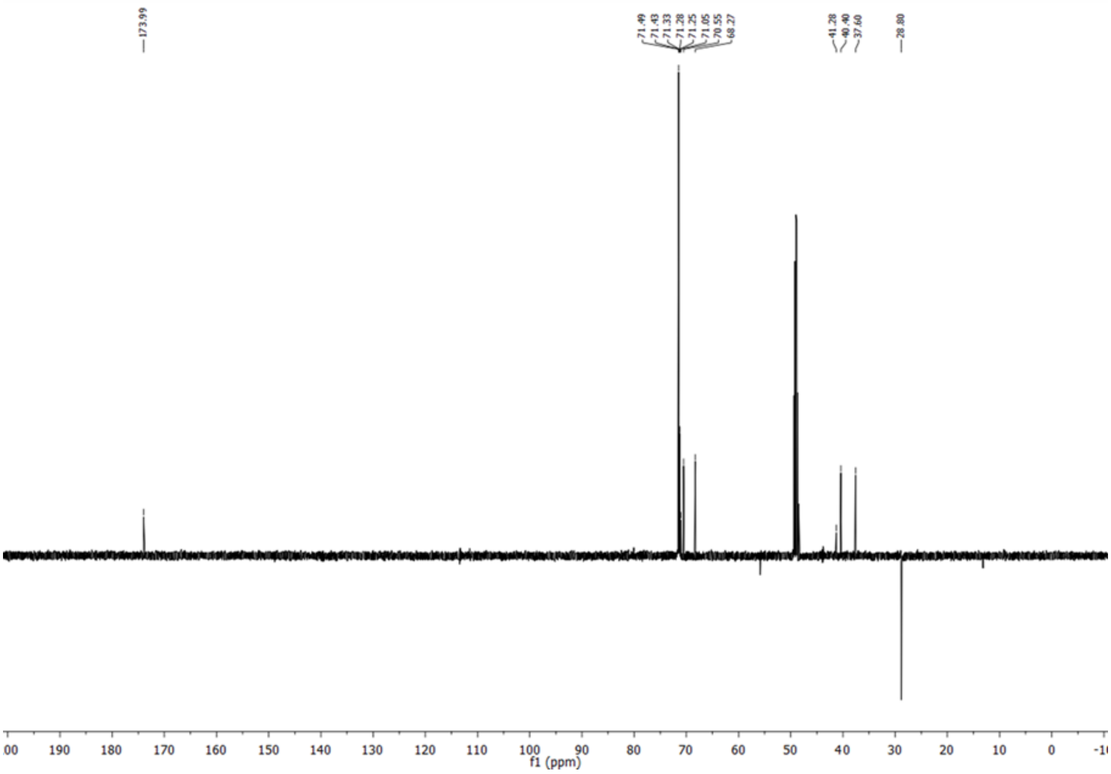

$^{1}H$ NMR (500 MHz, $CD_3OD$)

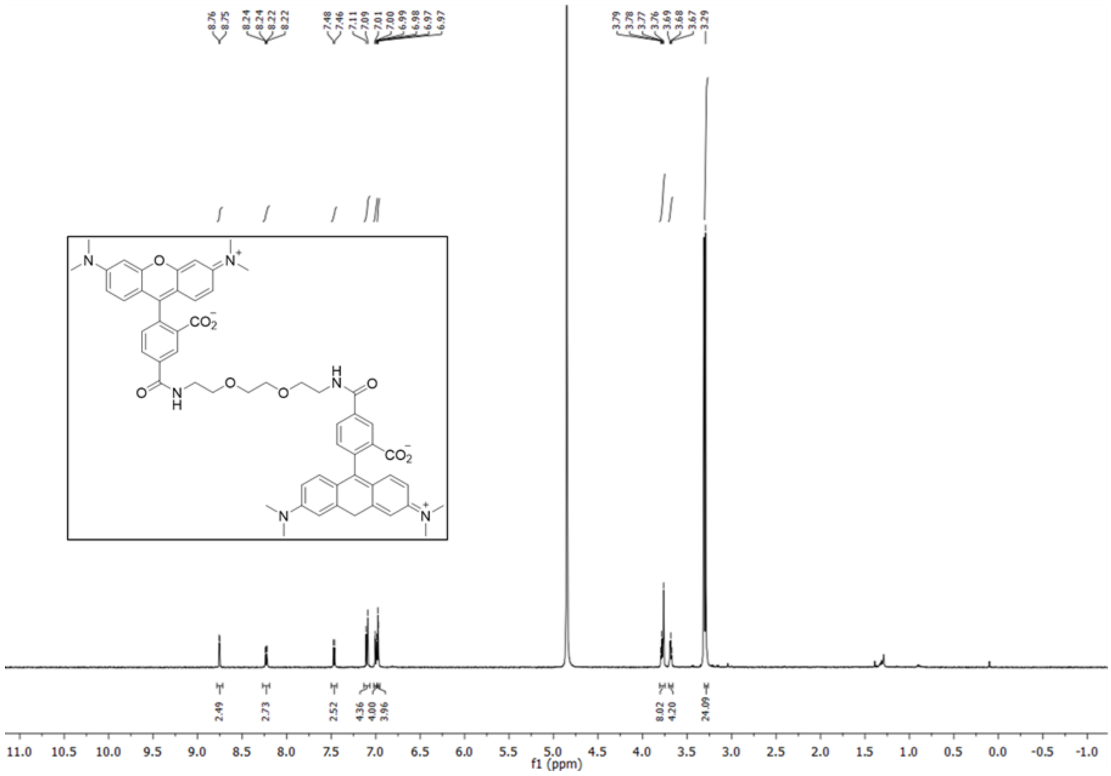

$^{13}C$ NMR (126 MHz, $CD_3OD$)

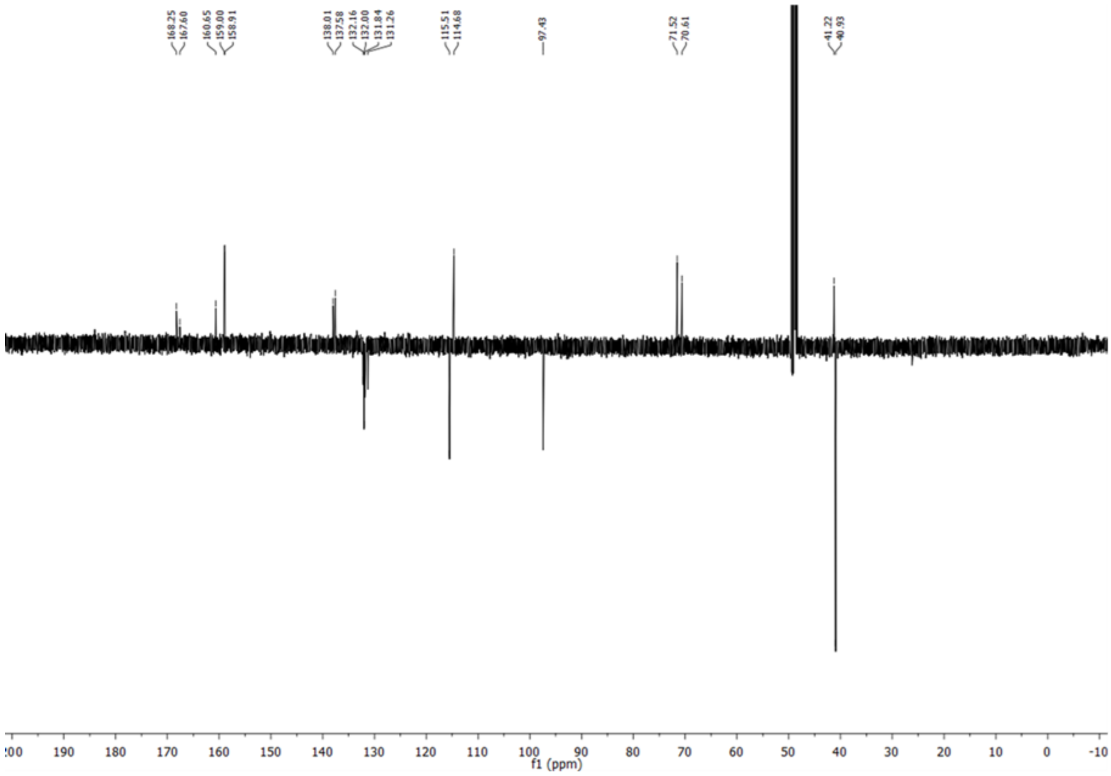

$^{1}H$ NMR (500 MHz, $CD_3OD$)

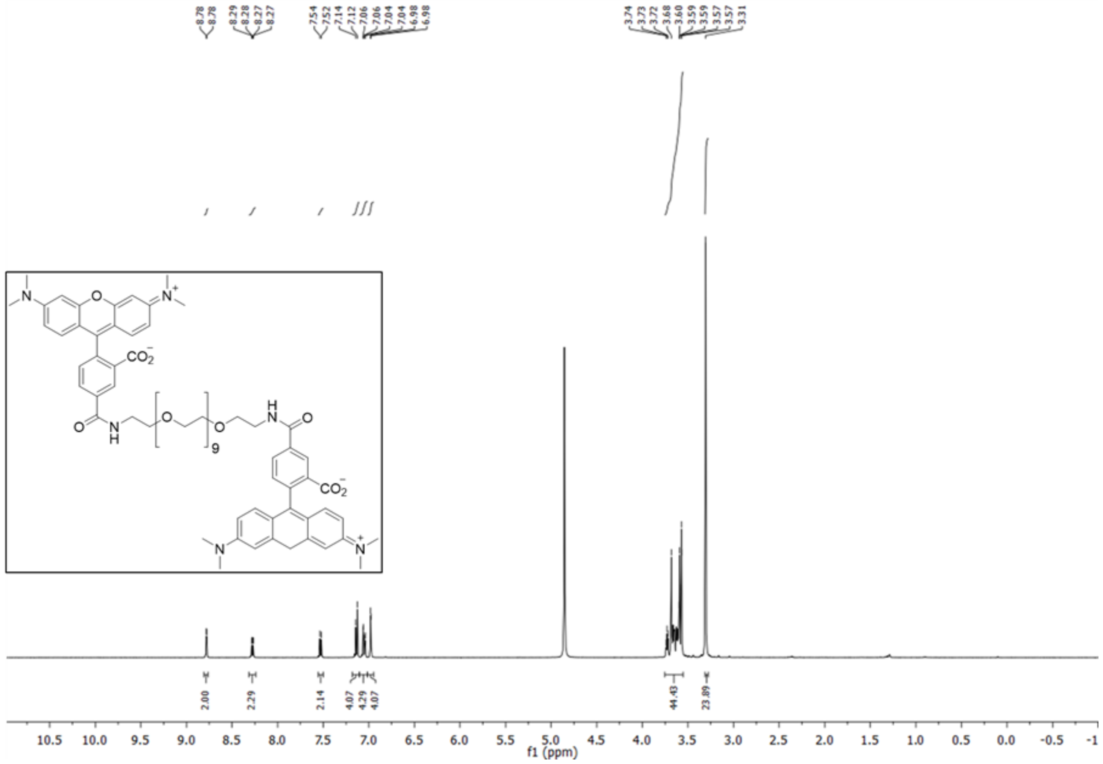

$^{13}C$ NMR (126 MHz, $CD_3OD$)

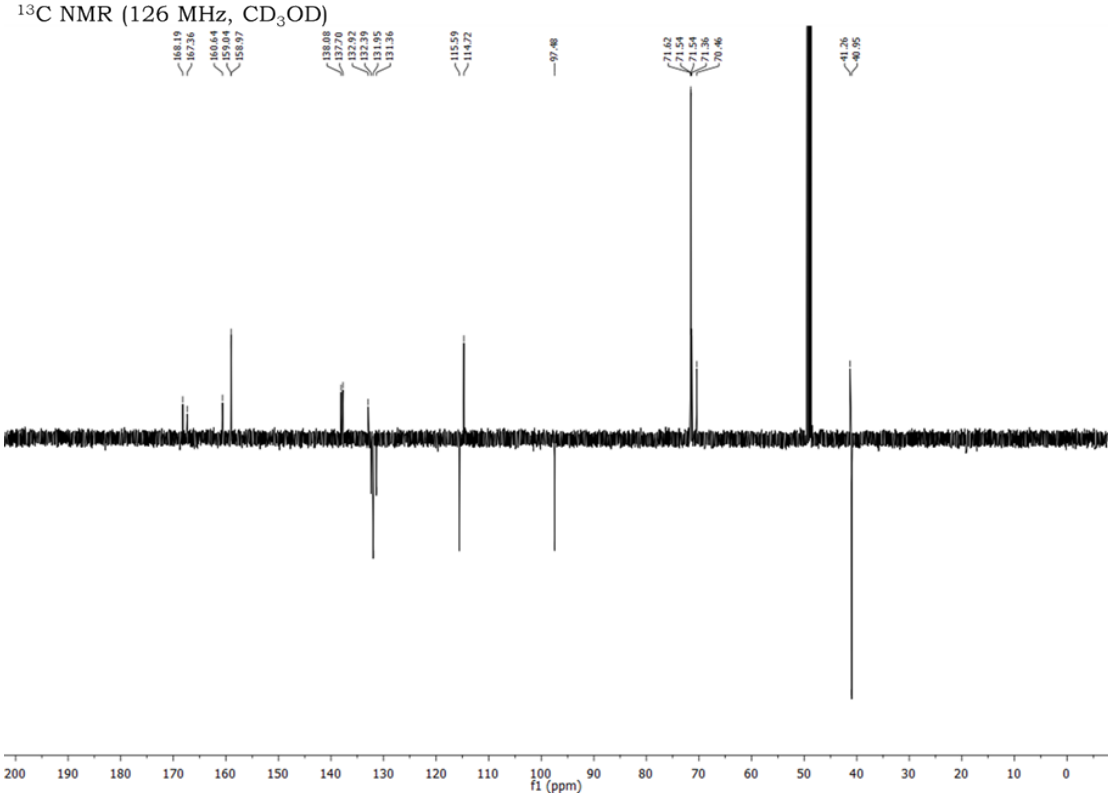

$^{1}H$ NMR (500 MHz, $CD_3OD$)

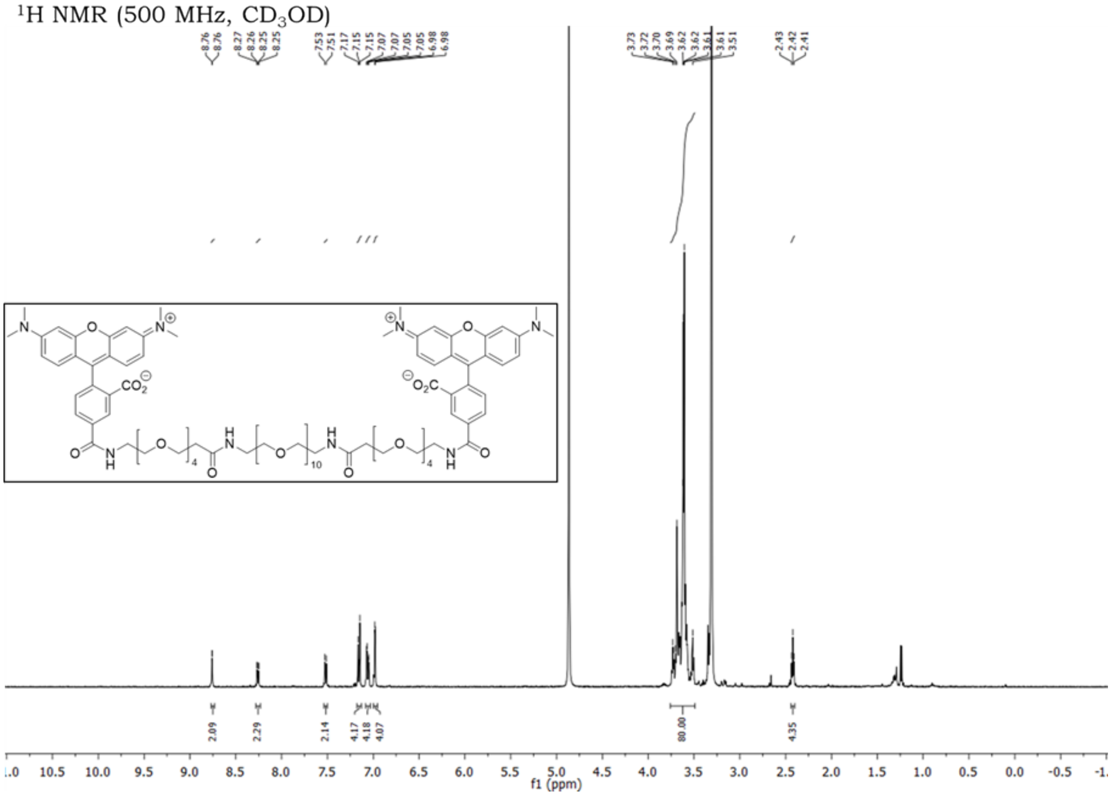


$^{13}C$ NMR (126 MHz, $CD_3OD$)

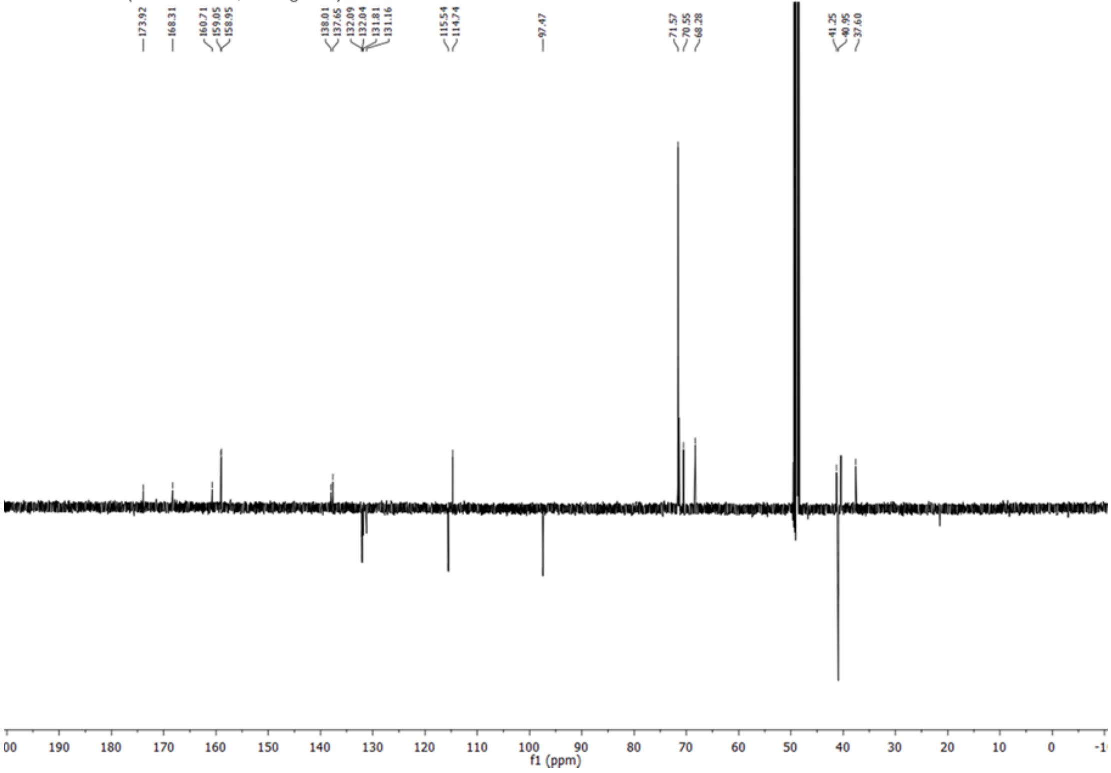

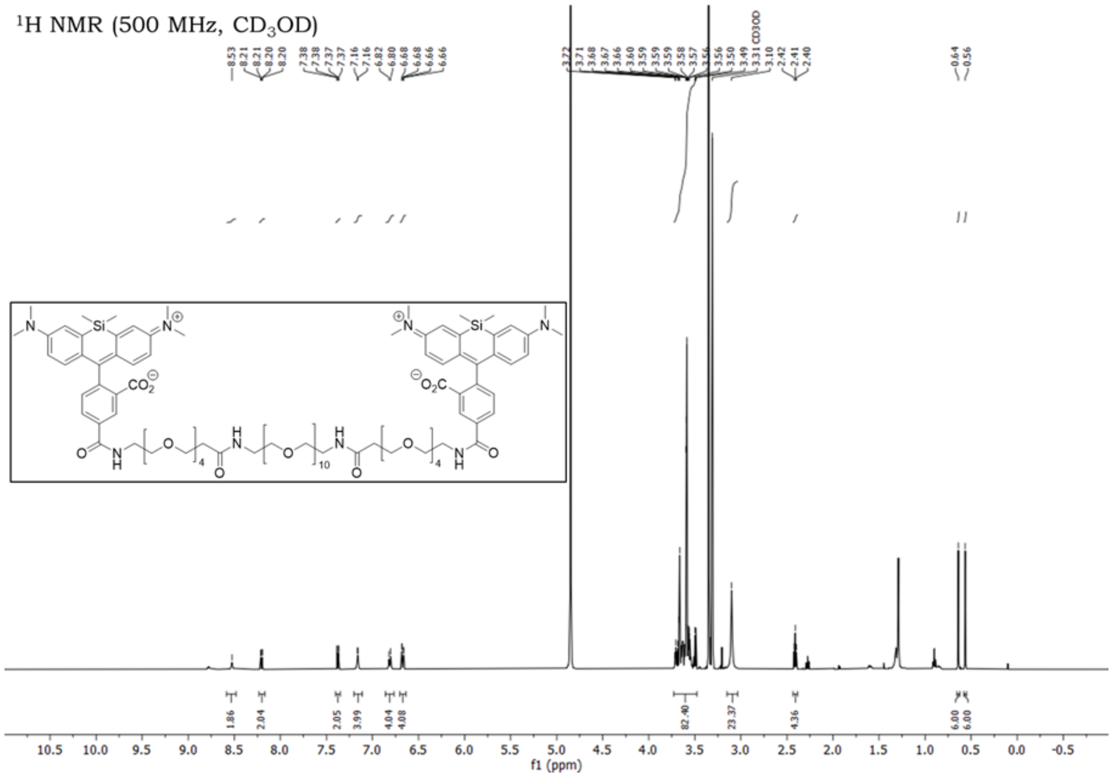

[1]H NMR (500 MHz, CD3OD)
f1 (ppm)

### Supplementary References


1. Terdale, S. & Tantray, A. Spectroscopic study of the dimerization of rhodamine 6G in water and different organic solvents. *J. Mol. Liq.* **225**, 662-671 (2017).
2. Bouhedda, F. et al. A dimerization-based fluorogenic dye-aptamer module for RNA imaging in live cells. *Nat. Chem. Biol.* **16**, 69-76 (2020).
3. Chapman, M. & Euler, W.B. Rhodamine 6G structural changes in water/ethanol mixed solvent. *J. Fluoresc.* **28**, 1431-1437 (2018).
4. Mack, E.T. et al. Dependence of avidity on linker length for a bivalent ligand–bivalent receptor model system. *J. Am. Chem. Soc.* **134**, 333-345 (2012).
5. Kitov, P.I. & Bundle, D.R. On the nature of the multivalency effect: a thermodynamic model. *J. Am. Chem. Soc.* **125**, 16271-16284 (2003).
6. Litke, J.L. & Jaffrey, S.R. Highly efficient expression of circular RNA aptamers in cells using autocatalytic transcripts. *Nat. Biotechnol.* **37**, 667-675 (2019).
7. Grubbs, R.D. Intracellular magnesium and magnesium buffering. *Biometals* **15**, 251-259 (2002).
8. Guo, J.U. & Bartel, D.P. RNA G-quadruplexes are globally unfolded in eukaryotic cells and depleted in bacteria. *Science* **353** (2016).
9. Chen, X. et al. Visualizing RNA dynamics in live cells with bright and stable fluorescent RNAs. *Nat. Biotechnol.* **37**, 1287-1293 (2019).
10. Li, X. et al. Fluorophore-Promoted RNA Folding and Photostability Enables Imaging of Single Broccoli-Tagged mRNAs in Live Mammalian Cells. *Angew. Chem. Int. Ed.* **59**, 4511-4518 (2020).
11. Cawte, A.D., Unrau, P.J. & Rueda, D.S. Live cell imaging of single RNA molecules with fluorogenic Mango II arrays. *Nat. Commun.* **11**, 1-11 (2020).
12. Braselmann, E. et al. A multicolor riboswitch-based platform for imaging of RNA in live mammalian cells. *Nat. Chem. Biol.* **14**, 964-971 (2018).
13. Wirth, R. et al. SiRA: a silicon rhodamine-binding aptamer for live-cell super-resolution RNA imaging. *J. Am. Chem. Soc.* **141**, 7562-7571 (2019).
14. Sunbul, M. et al. Super-resolution RNA imaging using a rhodamine-binding aptamer with fast exchange kinetics. *Nat. Biotechnol.* **39**, 686-690 (2021).
15. Bolte, S. & Cordelières, F.P. A guided tour into subcellular colocalization analysis in light microscopy. *J. Microsc.* **224**, 213-232 (2006).
16. Tarantino, N. et al. TNF and IL-1 exhibit distinct ubiquitin requirements for inducing NEMO–IKK supramolecular structures. *J. Cell Biol.* **204**, 231-245 (2014).
17. Fusco, D. et al. Single mRNA molecules demonstrate probabilistic movement in living mammalian cells. *Curr. Biol.* **13**, 161-167 (2003).
18. Trachman III, R.J. et al. Crystal structures of the Mango-II RNA aptamer reveal heterogeneous fluorophore binding and guide engineering of variants with improved selectivity and brightness. *Biochemistry* **57**, 3544-3548 (2018).
19. Tan, X. et al. Fluoromodules consisting of a promiscuous RNA aptamer and red or blue fluorogenic cyanine dyes: selection, characterization, and bioimaging. *J. Am. Chem. Soc.* **139**, 9001-9009 (2017).
20. Lukinavičius, G. et al. A near-infrared fluorophore for live-cell super-resolution microscopy of cellular proteins. *Nat. Chem.* **5**, 132-139 (2013).
21. McCann, T.E. et al. Activatable optical imaging with a silica-rhodamine based near infrared (SiR700) fluorophore: a comparison with cyanine based dyes. *Bioconjugate Chem.* **22**, 2531-2538 (2011).
22. Chiba, K. et al. Specific fluorescence labeling of target proteins by using a ligand–4-azidophthalimide conjugate. *Chem. Commun.* **53**, 8751-8754 (2017).
23. Robers, M.B. et al. A luminescent assay for real-time measurements of receptor endocytosis in living cells. *Anal. Biochem.* **489**, 1-8 (2015).
